# Evolutionarily diverged on-switch for actin assembly in fungal endocytosis

**DOI:** 10.1101/2025.06.22.660955

**Authors:** Juliette Sebellin, Fabina Binth Kandiyoth, Andrea Picco, Markku Hakala, Anne-Sophie Rivier-Cordey, Alphée Michelot, Christopher P. Toret, Marko Kaksonen

## Abstract

Clathrin-mediated endocytosis is a conserved eukaryotic trafficking process where an Arp2/3 complex nucleated branched actin network provides force for vesicle formation. The mechanisms that initiate endocytic actin assembly are incompletely understood. In the fission yeast, *Schizosaccharomyces pombe*, actin assembly is initiated by Dip1, an Arp2/3 activator. In the budding yeast, *Saccharomyces cerevisiae,* the initiation of actin assembly has remained a mystery. Here we show that *S. cerevisiae* Ldb17, the homolog of Dip1, functions as an on-switch for endocytic actin assembly. Unexpectedly, the regulation of Ldb17 is more complicated than that of constitutively active Dip1. Ldb17 is controlled by a coat protein, Sla1, via separate recruitment and activation steps. This regulation was likely lost in the *S. pombe* lineage and this simplification may be related to other changes in actin assembly between these species. Our findings add a key missing piece in the understanding of endocytosis in *S. cerevisiae* and reveal an intriguing evolutionary tinkering of the actin on-switch.

## INTRODUCTION

Clathrin-mediated endocytosis is a highly conserved eukaryotic process in which cells internalize plasma membrane vesicles for intracellular trafficking to regulate nutrient uptake, cell signaling, and membrane protein and lipid homeostasis (McMahon and Boucrot, 2011). The majority of the clathrin-mediated endocytic proteins are conserved in both fungi and animals (Dergai et al., 2016), and assemble sequentially at endocytic sites to temporally mediate endocytic event initiation, membrane bending and vesicle release (Kaksonen and Roux, 2018). However, the mechanisms of membrane bending and vesicle production have diverged between species. How this variation is achieved with largely conserved machinery remains unknown (Kaksonen and Roux, 2018). Even between the yeasts, *Saccharomyces cerevisiae* and *Schizosaccharomyces pombe*, a surprising endocytic variation occurs in protein amounts, sequential assemblies, and invagination and vesicle kinetics (Picco et al., 2024; Sun et al., 2019).

In fungi, actin is essential for membrane bending and vesicle formation, and actin assembly is precisely timed (∼10 seconds) during these final steps of the endocytic pathway (Picco et al., 2024; Sun et al., 2019). Specifically, the major endocytic Arp2/3 complex activators, WASP and class I myosin motors assemble around the endocytic coat. The coat contains Pan1, another Arp2/3 activator, and together these proteins stimulate assembly of Arp2/3-dependent branched actin networks at endocytic sites (Picco et al., 2015; Sun et al., 2006). These branched actin networks are anchored to the membrane and to the endocytic coat via Sla2, Ent1 and Ent2, clathrin-binding adaptor proteins. Actin network assembly at the plasma membrane propels the linked coat and associated membrane inward thereby driving vesicle budding (Kaksonen et al., 2003; Skruzny et al., 2012). In *S. cerevisiae,* WASP arrives ∼20s before actin assembly occurs (Picco et al., 2024; Sun et al., 2015). This pre-assembly of WASP is thought to facilitate an isotropic actin network assembly around the coat (Mund et al., 2018). However, in *S. pombe* WASP co-assembles with actin (Picco et al., 2024; Sirotkin et al., 2010, 2005). This shift in actin regulation and endocytic assembly divergence, is significant given the central role of actin in endocytosis, but how the actin assembly timing is controlled is unknown.

In *S. pombe*, Dip1 is a proposed trigger for actin assembly at endocytic sites (Basu and Chang, 2011). <u>W</u>ISH, <u>D</u>ip1 and <u>S</u>pin90 (WDS) proteins are an orthologous protein family that share a conserved structured core containing an Arp2/3 complex-binding region (Wagner et al., 2013). *Schizosaccharomyces pombe* Dip1 and mammalian Spin90 have a conserved function in actin filament nucleation (Cao et al., 2020; Kim et al., 2006; Wagner et al., 2013). Unlike other types of Arp2/3 activators, WDS proteins, like Dip1, are able to nucleate a new unbranched actin filament by activating the Arp2/3 complex independent of a preformed actin filament. Dip1 activates the Arp2/3 complex on a different face than WASP and does not depend on a WCA domain (Balzer et al., 2018; Wagner et al., 2013). The newly formed filament could provide the first filament for formation of an actin branch via WASP-dependent activation of the Arp2/3 complex at endocytic sites (Balzer et al., 2020). In the *S. pombe* endocytic pathway, WASP, Dip1 and actin all assemble at similar times and Dip1 may provide the initial switch that triggers WASP-activated branch assembly (Basu and Chang, 2011). WASP and Dip1 synergize their activities *in vitro*, which implies a potent cooperation for branched actin assembly at endocytic sites (Balzer et al., 2020). The Dip1 homologue in *S. cerevisiae*, Ldb17, localizes to endocytic sites just prior to actin assembly and mutants have endocytosis and actin patch defects (Burston et al., 2009), but has not been studied in detail in the budding yeast endocytic pathway.

The literature suggests a mechanistic role for Ldb17 and Dip1 in the nucleation of actin networks at endocytic sites. *S. cerevisiae* and *S. pombe* endocytic pathway assemblies differ in this critical moment of actin assembly. We sought to investigate WDS protein function at fungal endocytic sites to understand how Dip1 mechanisms relate to the *S. cerevisiae* pathway and uncover mechanistic divergences.

## RESULTS

### WDS proteins are very late components of the endocytic coat

We wanted to compare and contrast the assembly dynamics and motile behavior of WDS family proteins in three fungal species: *S. cerevisiae, S. pombe* and *Ustilago maydis*. These species have both described and diverged endocytic pathways and give phylogenetic context to understand mechanistic changes in endocytic processes (Picco et al., 2024). We tagged the WDS ortholog in each species endogenously at the C-terminus with mNeonGreen (mNG) in strains that also expressed an endogenous fimbrin ortholog tagged with mCherry (mCh). Fimbrin marks the actin filaments at endocytic sites and provides a temporal landmark for endocytic actin-dependent vesicle formation stage (Kaksonen et al., 2005; Sirotkin et al., 2010). These cells were imaged at the mid plane with live epifluorescence microscopy. In all three species, the WDS family proteins localized to cortical patches that partially colocalized with fimbrin patches. This observation suggested that the endocytic role of the WDS proteins is broadly conserved in dikarya fungi (Figure 1A). WDS protein signals were weak in all species, therefore we used TIRF microscopy to minimize photobleaching and cytoplasmic background to analyze the timing of WDS protein localization at endocytic sites. In *S.cerevisiae*, Ldb17 arrived less than 5 s prior to actin assembly at endocytic sites, which is consistent with prior observations (Burston et al., 2009). In comparison, *S. pombe* (Dip1) and *U. maydis* (UMAG_01405, henceforth called Dip1) orthologs appeared just before fimbrin (Figure 1B), and in agreement with earlier observations, which showed that *S.pombe* Dip1 and Wsp1 arrive ∼1 s before actin assembly (Basu and Chang, 2011). We found that Ldb17 had a slightly longer lifetime (∼10 s) than the other two Dip1s, ∼9 s (*S. pombe* and *U. maydis*) (Figure 1C), which is consistent with the recruitment differences between species. These results suggest a similar arrival for WDS proteins just prior to actin assembly, which is well conserved in dikarya and consistent with an actin trigger function.

**Figure 1:**
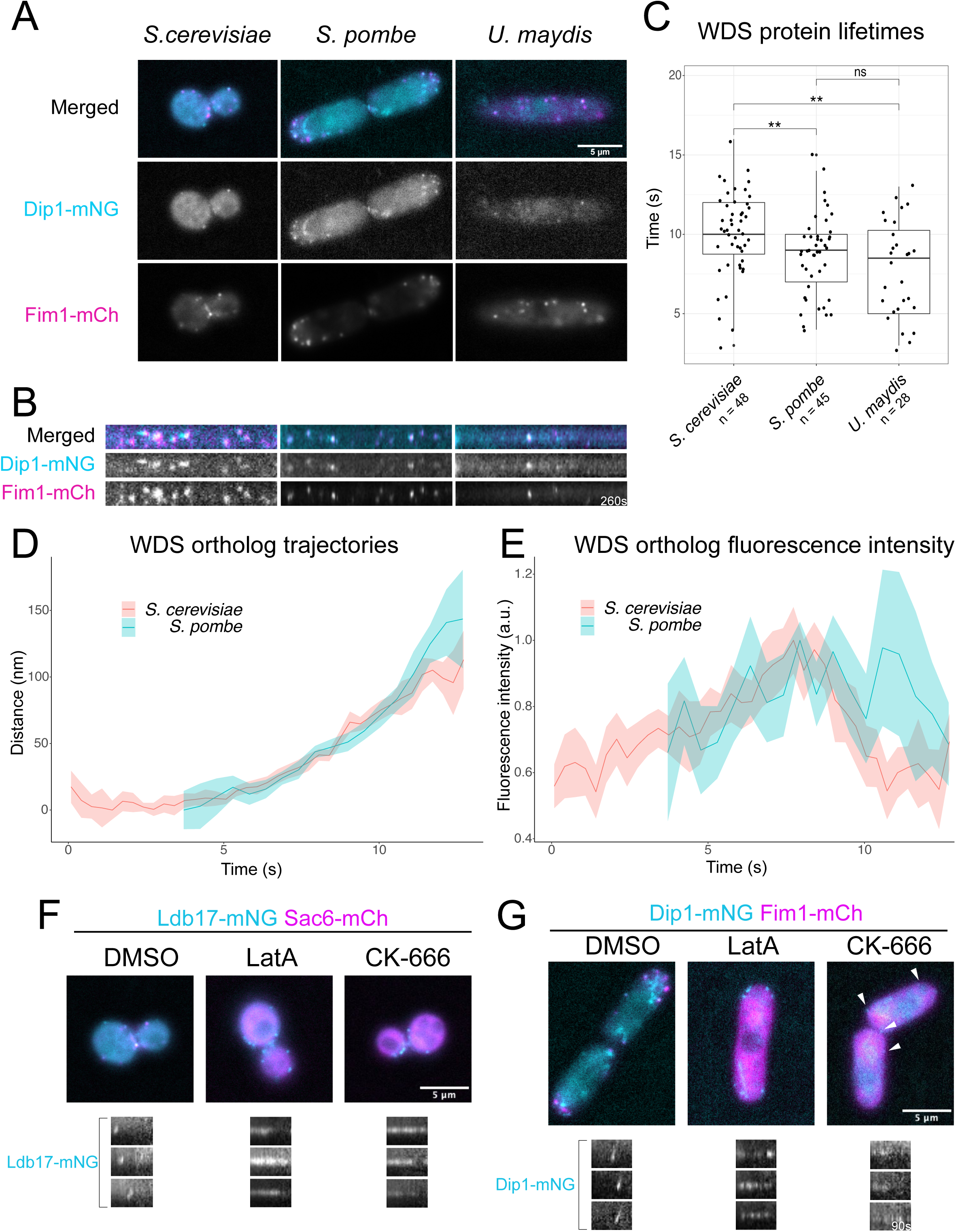
WDS protein localization. (A) Equatorial epifluorescence images for WDS and Fim1 orthologs in each species. (B) Kymograph of 2-color TIRF microscopy movies that show indicated WDS ortholog (cyan) and co-expressed Fim1 ortholog (magenta) in each of the three species. (C) Boxplot showing WDS orthologs lifetime. (D) Trajectory plot of WDS orthologs in *S.cerevisiae* (red) and *S. pombe* (blue). Error is the 95% confidence interval. (E) Fluorescence intensity profile of WDS orthologs in *S.cerevisiae* (red) and *S. pombe* (blue). Error is the 95% confidence interval. (F) Equatorial epifluorescence images of Ldb17(cyan) and Sac6 (magenta) treated with DMSO, LatA and CK-666. (G) Equatorial epifluorescence images of Dip1 (cyan) and Fim1 (magenta) treated with DMSO, LatA and CK-666. Arrows in white.

Since the WASP assembly mechanisms diverged in *S. cerevisiae* and *S. pombe* (Picco et al., 2024), we focused on these two species. We imaged Ldb17 and Dip1 patches at equatorial planes in *S. cerevisiae* and in *S. pombe*, respectively, and then applied particle tracking of the endocytic patches. Particle tracking provides information about the relative position and movement of endocytic proteins during vesicle formation. Both orthologs internalized at the end of the patch lifetime at similar speeds for ∼100-150 nm (Figure 1D). Ldb17 fluorescence intensity peaked at 7s before falling precipitously. Dip1 showed a similar pattern of assembly and disassembly, but the signal was noisier due to a weaker signal at endocytic sites in *S. pombe*, likely due to lower number of molecules compared to *S. cerevisiae* (Figure 1E). The motile behaviors of Ldb17 and Dip1 protein are consistent with endocytic coat proteins that internalize at a similar initial rate in *S. pombe* and *S. cerevisiae* (Kaksonen et al., 2005; Sirotkin et al., 2010). The motile behavior is also consistent with Dip1 in binding to and activating the Arp2/3 complex to form the first actin filaments at the endocytic site. These first filaments would be bound to coat proteins and would internalize with the coat as the actin network continues to grow from the surface of the plasma membrane (Wagner et al., 2013b).

Both Dip1 and Ldb17 localize independent of actin (Basu and Chang, 2011; Burston et al., 2009). We further analyzed the dynamics of these proteins by inhibiting actin assembly with Latrunculin A (LatA) or CK-666. LatA sequesters actin monomers and depolymerizes actin filaments (Ayscough et al., 1997) whereas CK-666 is an Arp2/3 inhibitor (Nolen et al., 2009). In the LatA-treated *S. cerevisiae* cells, persistent Ldb17 patches transiently flickered at the membrane for the duration of the movie (2.5 min) and remained immotile (Figure 1F). Dip1 patches similarly flickered and remained immotile in LatA-treated *S.pombe* cells (Figure 1G). In the presence of CK-666, Ldb17 and Dip1 were both recruited to persistent, nonmotile patches that flickered similar to LatA (Figure 1F and 1G). However, unlike Ldb17, Dip1 recruitment was weaker in the presence of CK-666 relative to LatA (Figure 1G, white arrowheads), which suggests the Arp2/3-inhibition has a stronger effect on Dip1 behavior. Together, these results indicate that WDS proteins are recruited independently of actin, while Dip1 has an additional sensitivity for Arp2/3 function and hints at a potential difference between orthologs. Moreover, actin dynamics are essential for WDS protein movement and normal disassembly at endocytic sites, which suggested a largely shared relationship with the actin cytoskeleton.

### Dip1 and Ldb17: similar mutant phenotypes but different actin biochemistries

We next compared WDS protein function in the two species by deletion of the genes. Previously, *S. pombe dip1* deletion was shown to delay the initiation of actin assembly at endocytic sites (Basu and Chang, 2011). We imaged endocytic proteins Sla1-EGFP as a marker for the vesicle coat and Sac6-mCh as a marker for endocytic actin in *S. cerevisiae* and their respective orthologs in *S. pombe*, Shd1-sfGFP and Fim1-mCherry. In wildtype *S. cerevisiae* cells, Sla1 first appeared at the plasma membrane as immotile patches and then internalized with actin assembly (Figure 2A) as previously described (Kaksonen et al., 2003). In Ldb17 null cells, Sla1-EGFP patches still internalized with the arrival of actin similar to WT (Figure 2A). However, a marked difference occurred in the early immotile phase, which was dramatically extended (Figure 2A, kymographs). In *S. pombe,* Dip1 deletion resulted in a similar extension of the immotile phase of Shd1-sfGFP patches prior to actin-associated internalization (Figure 2B). In addition, in *S. pombe,* Dip1 mutant cells were shorter and rounder than WT cells, while no obvious cell shape change was seen in *S. cerevisiae* (Figure 2A and B). These results suggest a conserved role for the WDS proteins in regulating endocytosis in these two species.

**Figure 2:**
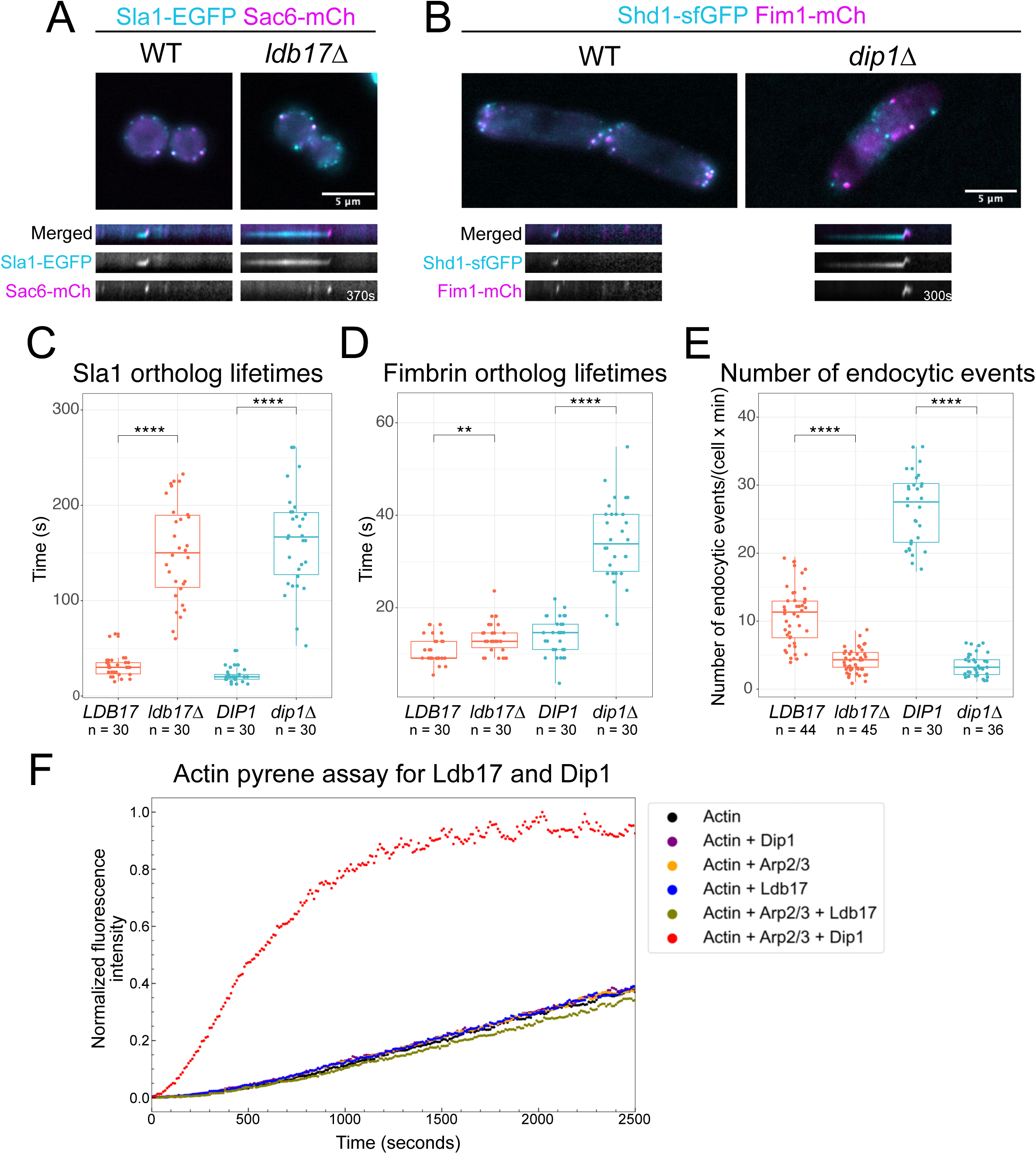
WDS protein and endocytic protein dynamics. (A) Equatorial epifluorescence images of *S.cerevisiae* for the coat marker Sla1-EGFP (cyan) and the actin marker Sac6-mCherry (magenta) in each species in WT and Ldb17 knock-out. Associated kymographs are displayed below. (B) Equatorial epifluorescence images of *S. pombe* for the coat marker Shd1-EGFP (cyan) and the actin marker Fim1-mCherry (magenta) in each species in WT and Ldb17 knock-out. Associated kymographs are displayed below. (C) Boxplot of coat lifetime in *S.cerevisiae* and *S. pombe* using Sla1-EGFP or Shd1-EGFP in the context of *ldb17Δ*, *dip1Δ* and their respective WT.(D) Boxplot of actin lifetime in *S.cerevisiae* and *S. pombe* using Sac6-mCherry or Fim1-mCherry in the context of *ldb17Δ*, *dip1Δ* and their respective WT. (E) Number of endocytic events per cell per minute in the context of *ldb17Δ*, *dip1Δ* and their respective WT. (F) Time course of actin polymerization monitored by pyrene fluorescence. The protein concentrations are 2 µM for actin, 0.1 µM for Arp2/3 complex and 1 µM for all the other proteins.

The lifetimes for Sla1 patches significantly extended and were more variable in both *ldb17Δ* and *dip1Δ* mutants relative to the wildtypes (Figure 2C). Fimbrin patch lifetimes were also extended in the mutants in both species but the effect was much more dramatic in *S. pombe* compared to *S. cerevisiae* (Figure 2D). We then counted the number of Sla1-EGFP patches that internalized per cell within one minute. This analysis showed a significant reduction in the frequency of endocytic events in the WDS protein mutants (Figure 2E). These results reveal a similar delay for both species in actin assembly at endocytic sites in WDS protein mutants.

The similarity of the deletion phenotypes suggests that *S. cerevisiae* Ldb17 functions similarly to *S. pombe* Dip1 by directly activating the Arp2/3 complex to generate the first actin filaments at the endocytic site (Wagner et al., 2013). To test this idea, we expressed and purified both Ldb17 and Dip1 to compare their Arp2/3 activation activity with pyrene actin assay with purified *S.cerevisiae* actin and Arp2/3 complex. Addition of Ldb17 or Dip1 with actin alone resulted in no measurable effect on actin assembly (Figure 2F). Dip1 resulted in strong activation of Arp2/3-mediated actin polymerization (Figure 2F), which is consistent with earlier experiments that used Dip1 with *S. pombe* Arp2/3 proteins (Balzer et al., 2020; Wagner et al., 2013). In contrast, Ldb17 had little to no effect on actin polymerization rate when added to actin and Arp2/3 (Figure 2F). This contrasting result suggests that despite conserved localizations and deletion phenotypes, there is a striking difference in the mechanism of Arp2/3 activation between Ldb17 and Dip1. We speculated that Ldb17 may require an additional factor for its activation in contrast to Dip1.

### Sequence comparison suggests evolution in the regulation of WDS proteins

The biochemical difference between Ldb17 and Dip1 indicates that these proteins may differ in the way their activities are regulated. To better understand the molecular basis of this difference and its evolution we compared sequences of WDS protein homologs from a range of different organisms. Homology searches revealed that WDS family proteins are present in the supergroup of amorphea, which contains amoebozoa, apusozoa, holozoa, which includes animals, and holomycota, which includes fungi. We focused on these groups and identified sequences of WDS orthologs from several species across these clades (Supplemental text). We then aligned the WDS protein sequences and compared their features. WDS proteins share a homologous central core region, which has been shown to be composed of alpha helical repeats (Figure 3A) (Luan et al., 2018; Shaaban et al., 2020). In Dip1 and SPIN90 the C-terminal portion of this core region was demonstrated to bind and activate the Arp2/3 complex (Luan et al., 2018; Shaaban et al., 2020). This C-terminal portion has the highest degree sequence conservation among the orthologs (Figure S1A) (Liu et al., 2022). In comparison, the N-terminal half of the core region diverges greater in sequence (Figure 3A and Figure S1A). The function of the N-terminal part of the core is not as well characterized, but may interact with regulatory proteins (Luan et al., 2018). Proteins that have high similarity to a WDS core were found in all amorphea species that we searched. However, additional sequence features were found in opisthokonta species (holozoans and holomycota). The orthologs in the holozoan branch, like Spin90, have an N-terminal extension that often harbor an SH3 domain and also often contain proline-rich regions (PRR) and mediates several Spin90 interactions (Cao et al., 2020; Kim et al., 2006b; Lim et al., 2003). SH3 domains are a common feature of actin and endocytic proteins and are established PRR interaction domains (Hummel and Kaksonen, 2023; Tonikian et al., 2009). In contrast to holozoa, orthologs in the holomycotan branch contain a C-terminal extension that includes a PRR that is often flanked by an arginine or lysine charge-rich patch (Figure 3B and Supplemental text). In fungi, the C-terminal extension was independently lost in some unique branches (Figure 3B). The *S. pombe* ortholog represents such a lineage (Figure 3B). Recently, an N-terminal portion of animal Spin90s were shown to dimerize and generate bidirectional actin filaments, which represents another divergence with holomycota (Liu et al., 2025).

**Figure 3:**
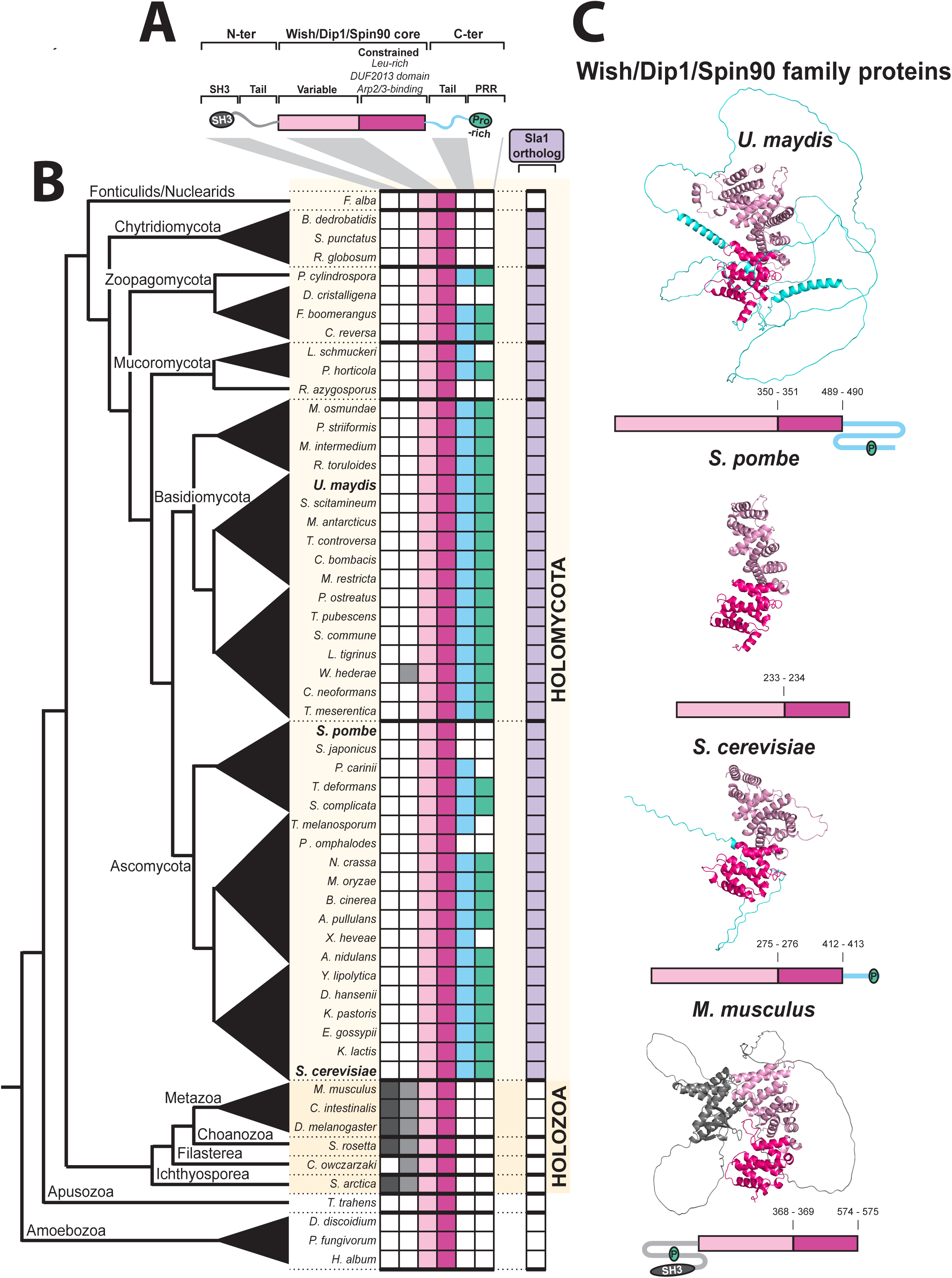
WDS protein family. (A) Top schematic shows generalized WDS ortholog organization. For each species a dot indicates presence of N-terminal tail, N-terminal SH3 domain, variable core, constrained core, C-terminal PRR or C-terminal tail. (B) Selected WDS orthologs in amorphea. Species and accession numbers are indicated and are organized based on relative position on the tree of life (left). Boxes indicate the presence of regions depicted in A. Right most column indicates the presence of a Sla1 ortholog in the species. (C) Alpha fold structures of WDS orthologs of indicated species. WDS core regions variable (light blue) and constrained (dark blue) and tails are (color). All orthologs cores were aligned and oriented in the same direction in PyMOL

The structures of the C-terminal core regions of *S. pombe* Dip1 and the whole core for mammalian Spin90 were experimentally solved (Liu et al., 2022; Luan et al., 2018). We generated AlphaFoldV2.0 structure predictions for the orthologs in *S. cerevisiae*, *S.pombe*, *Ustilago maydis* and *Mus musculus* (Figure 3C). The structure prediction of *S. pombe* Dip1 conformed well to the published experimental structure (Figure S1B). The AlphaFold predictions suggested a structured WDS core for each ortholog (Figure 3C and S1). The predictions also suggested that the opisthokonta extensions, whether N-ter or C-ter, are largely unstructured apart from a few potential helical regions and the SH3 domain in *Mus musculus*. This analysis suggests that the ancestral form of WDS proteins was likely composed of only the structured core. In opisthokonts the WDS proteins gained N- or C-terminal regions, which may regulate activity or recruitment of the protein, likely involving PRR-SH3 interactions. The Dip1 protein of *S. pombe* represents a reversion to pre-opisthokonta-like core-only state. These sequence changes may contribute to the functional differences between Dip1 and Ldb17.

### Ldb17 regulation of the endocytic site is associated with Sla1

We next focused on C-terminally-tailed Ldb17 to better understand its function at the endocytic site. The *ldb17Δ* resulted in strongly extended and variable lifetime of the late coat phase followed by relatively normal actin patch assembly and coat internalization (Figure 2). This phenotype is remarkably similar to the phenotype reported for *sla1Δ* mutant (Kaksonen et al., 2005). Furthermore, Sla1 and Ldb17 have been reported to directly interact via Sla1’s SH3 domains (Burston et al., 2009). To investigate a possible functional relationship between Ldb17 and Sla1, we analyzed the assembly dynamics of another late coat protein, Pan1, and Sac6 in *ldb17Δ*, *sla1Δ*, and *ldb17Δ sla1Δ* mutants. As expected, Pan1-EGFP behaved similarly to our previous Sla1-EGFP observations in *ldb17Δ* and WT cells (Figure 2A and 4A). In *sla1Δ* cells, Pan1-EGFP patch lifetimes were extended and highly variable in duration. The Pan1-EGFP patches disappeared with the arrival of Sac6 (Figure 4A), as previously described (Kaksonen et al., 2005). When both Ldb17 and Sla1 were deleted the Pan1-EGFP phenotype appeared equivalent to single deletions (Figure 4A). We quantified Pan1 and Sac6 patch lifetimes and the frequency of endocytic events. These data showed that Pan1 patch lifetime in all three mutants was significantly longer and more variable than in WT (Figure 4B). These results suggest that both Ldb17 and Sla1 regulate the initiation of actin assembly. Fimbrin lifetimes were extended in all three mutants relative to WT, but the double mutant showed an exacerbated phenotype (Figure 4C). These data suggest an additive effect on actin patch lifetime, which may not be directly related to the initiation of actin assembly. The number of endocytic events per cell per minute was significantly reduced in all three mutants compared to the WT (Figure 4D), which revealed that all three mutants decrease the endocytic efficiency. We then quantified Pan1 patch internalization trajectories in each strain. In WT and *ldb17Δ* cells, Pan1 internalized approximately 150 nm and the internalization trajectories were indistinguishable (Figure 4E). In *sla1Δ* and *ldb17Δ sla1Δ* mutants the internalization trajectories differed from WT. These trajectories internalized a distance of ∼300 nm (Figure 4E). These data suggest that Sla1 likely also has functions that are not shared with Ldb17, which are related to internalization of the coat module, but not timing of the initiation. For example, Sla1 negatively regulates Las17 (Rodal et al., 2003).

**Figure 4:**
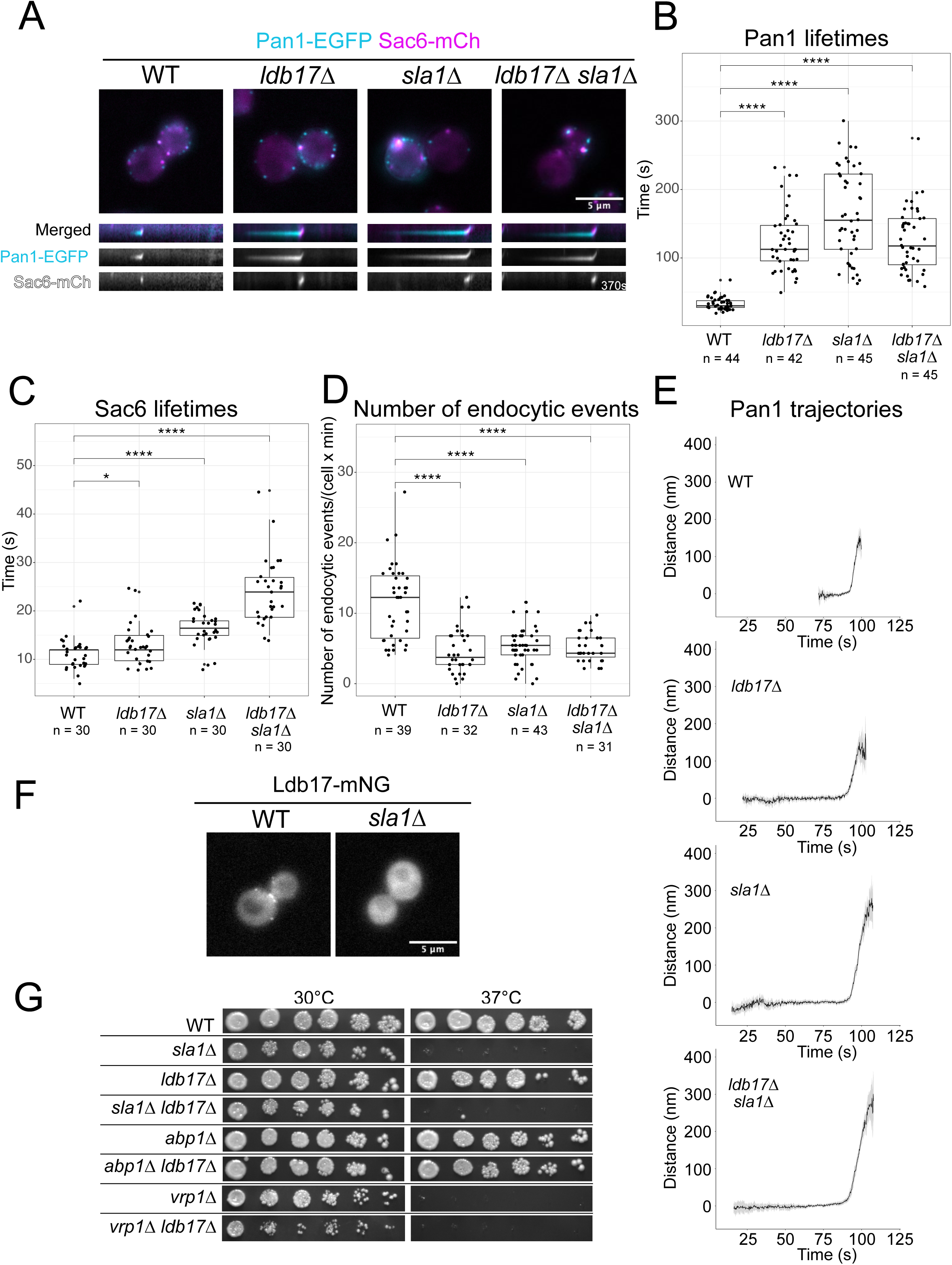
WDS protein function at endocytic sites. (A) *S.cerevisiae* equatorial epifluorescence images for the coat marker Pan1-EGFP, the actin marker Sac6-mCherry in WT, *ldb17Δ*, *sla1Δ*, and *ldb17Δ sla1Δ* cell backgrounds, and associated kymographs. (B) Boxplot of Pan1-EGFP lifetime in WT, *ldb17Δ*, *sla1Δ*, and *ldb17Δ sla1Δ* cell backgrounds. Pan1 lifetime difference is significantly different between *ldb17Δ* and *sla1Δ* and between *sla1Δ* and *ldb17Δ sla1Δ* at a two stars level. Pan1 lifetime difference between *ldb17Δ* and *ldb17Δ sla1Δ* is not significant. (C) Boxplot of Sac6-mCherry lifetime in WT, *ldb17Δ*, *sla1Δ*, and *ldb17Δ sla1Δ* cell backgrounds. (D) Boxplot of the number of endocytic events per cell per minute in WT, *ldb17Δ*, *sla1Δ*, and *ldb17Δ sla1Δ* cell backgrounds. (E) Trajectory plots of Pan1-EGFP in WT, *ldb17Δ*, *sla1Δ*, and *ldb17Δ sla1Δ* cells. Error is the 95% confidence interval (F) Upper two panels: Equatorial epifluorescence images of Ldb17-mNG in WT and *sla1Δ*. Bottom two panels: Equatorial epifluorescence images of Dip1-mNG in WT and *shd1Δ* cells. Dip1 patches are shown with white arrowheads (G) Yeast growth assay of eight strains conducted over 2 days at 30 and 37°C.

We tested the Sla1-dependency of Ldb17 recruitment to patches. In *sla1Δ* cells, Ldb17-mNG patches were not detected (Figure 4F). In contrast, Sla1 patch recruitment occurred in *ldb17Δ* cells (Figure 4E). In agreement with earlier observations (Burston et al., 2009), these results suggest that Sla1 acts upstream of Ldb17 protein recruitment and is consistent with the recruitment time of each protein to the endocytic site (Ldb17 arrives after Sla1). In conclusion, the *sla1Δ* phenotype, particularly the actin initiation delay, is likely due to the failed recruitment of Ldb17 to the patch.

To further understand Sla1-Ldb17 functions we analyzed genetic interactions. Mutant *sla1Δ* cells have reported genetic synthetic phenotypes with *abp1Δ* and *vrp1Δ* (Costanzo et al., 2010; Holtzman et al., 1993a). In contrast, *ldb17Δ* mutant was reported to suppress *vrp1Δ* phenotypes in systematic studies (Costanzo et al., 2016). To investigate these interactions we performed yeast growth assays of different mutant combinations at 30 and 37°C. As previously reported (Holtzman et al., 1993b), *sla1Δ* mutants had a growth defect at 37°C (Figure 4G), while *ldb17Δ* had no detectable growth defect relative to WT at any temperature tested (Figure 4G). The *sla1Δ* mutant growth phenotype was epistatic in regards to *ldb17Δ* (Figure 4G). The *ldb17Δ abp1Δ* mutant grew similar to WT. In contrast to the systematic data, the phenotypic suppression of *vrp1Δ* with *ldb17Δ* was not reproduced in our conditions, but we observed that *ldb17Δ vrp1Δ* cells had a more severe growth defect than *vrp1Δ* cells (Figure 4G). This genetic exacerbation of *vrp1Δ* phenotypes is similar for both *ldb17Δ* and *sla1Δ* mutants. Together these results suggest that Sla1 has both Ldb17-dependent and independent functions.

### The assembly dynamics of Las17 are altered in *ldb17Δ* cells

In *S. cerevisiae*, Las17 (WASP) assembles at endocytic sites about 20 seconds prior to actin assembly starts. This Las17 pre-assembly is partially mediated by Sla1 (Sun et al., 2017). Sla1 has also been shown to inhibit Las17 nucleation promoting activity *in vitro* (Rodal et al., 2003). This link is thought to play a role in regulating the timing of actin assembly and WASP concentration at the endocytic site (Sun et al., 2017). We investigated Las17 assembly dynamics at endocytic sites in relation to Ldb17 function. We analyzed Las17-EGFP in *ldb17Δ*, *sla1Δ* and *ldb17Δ sla1Δ* cells that co-expressed Sac6-mCherry. In WT cells, Las17-EGFP pre-assembles at endocytic sites and disappears after actin assembly (Figure 5A; Kaksonen et al., 2003). In *ldb17Δ* cells, *sla1Δ* cells and *ldb17Δ sla1Δ* cells, the time that Las17 resided at the membrane prior to actin assembly was dramatically increased compared to the WT cells (Figure 5A). Quantification revealed that in WT cells Las17 started to pre-assemble ∼20 seconds before the start of actin assembly. In all three mutants, the pre-assembly period was extended and highly variable with an average over 100s (Figure 5B). These observations are consistent with the extended lifetimes of Pan1 and Sla1 (Figures 2C and 4B) and indicate that the normal timing of actin assembly is impaired in *ldb17Δ* and *sla1Δ* mutants. We next quantitatively analyzed the fluorescence intensity of Las17-EGFP and Sac6-mCherry over time for WT and the mutants. In WT cells, Las17 intensity steadily ascends and peaks with actin assembly, before rapidly disappearing (Figure 5C). In *ldb17Δ*, *sla1Δ* and *ldb17Δ sla1Δ* cells, the ascending phase occurred throughout the extended pre-assembly phase, (Figure 5C). In *ldb17Δ* cells, like WT, the ascending phase was continuous and reached its maximum with actin polymerization. In contrast, in *sla1Δ* and *ldb17Δ sla1Δ* cells, had a steady ascending phase, which was followed by a rapid intensity spike when actin assembly appeared (Figure 5C). These results suggest that Sla1 has at least two activities on Las17. One regulates the timing of actin assembly, likely via Ldb17, and a second activity contributes to the pre-assembly of Las17 prior to actin assembly.

**Figure 5:**
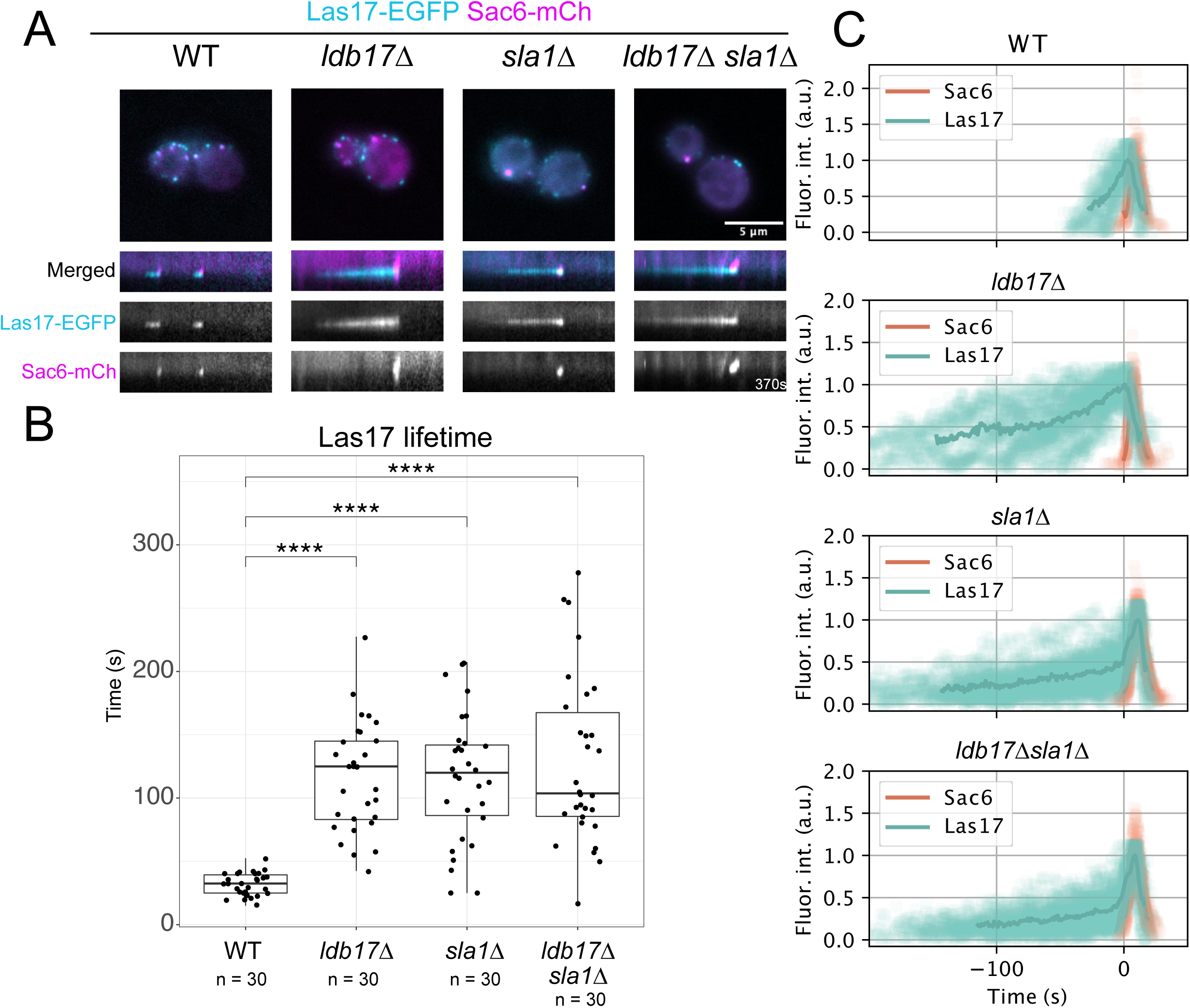
WDS protein and WASP dynamics. (A) *S.cerevisiae* equatorial epifluorescence images for the WASP protein Las17-EGFP, the actin marker Sac6-mCherry in WT, *ldb17Δ*, *sla1Δ*, and *ldb17Δ sla1Δ* cell backgrounds, and associated kymographs. (B) Boxplot of Las17-EGFP lifetime in WT, *ldb17Δ*, *sla1Δ*, and *ldb17Δ sla1Δ* cell backgrounds. The comparison of the lifetime in the three mutants showed no significant difference. (C) Fluorescence intensity profiles of Las17-EGFP aligned in respect to the corresponding Sac6-mCherry peaks in WT, *ldb17Δ*, *sla1Δ*, and *ldb17Δ sla1Δ* cell backgrounds.

### Ldb17 regulation via SH3 proteins Bzz1 and Sla1

We next sought to better understand Ldb17 regulation and recruitment to endocytic sites. The PRR in the unstructured tail of Ldb17 (Figure 3) suggests a possible link with SH3 domain proteins, which could function as regulators or recruiters for Ldb17. We selected four candidate proteins for Ldb17 interactions: Lsb3 and Lsb4 (genome-duplicated orthologs), Bzz1, and Sla1 (Figure 6A), which have reported WDS protein interactions and arrive at endocytic sites before Ldb17 (Burston et al., 2009; Kim et al., 2006a; Tonikian et al., 2009).

**Figure 6:**
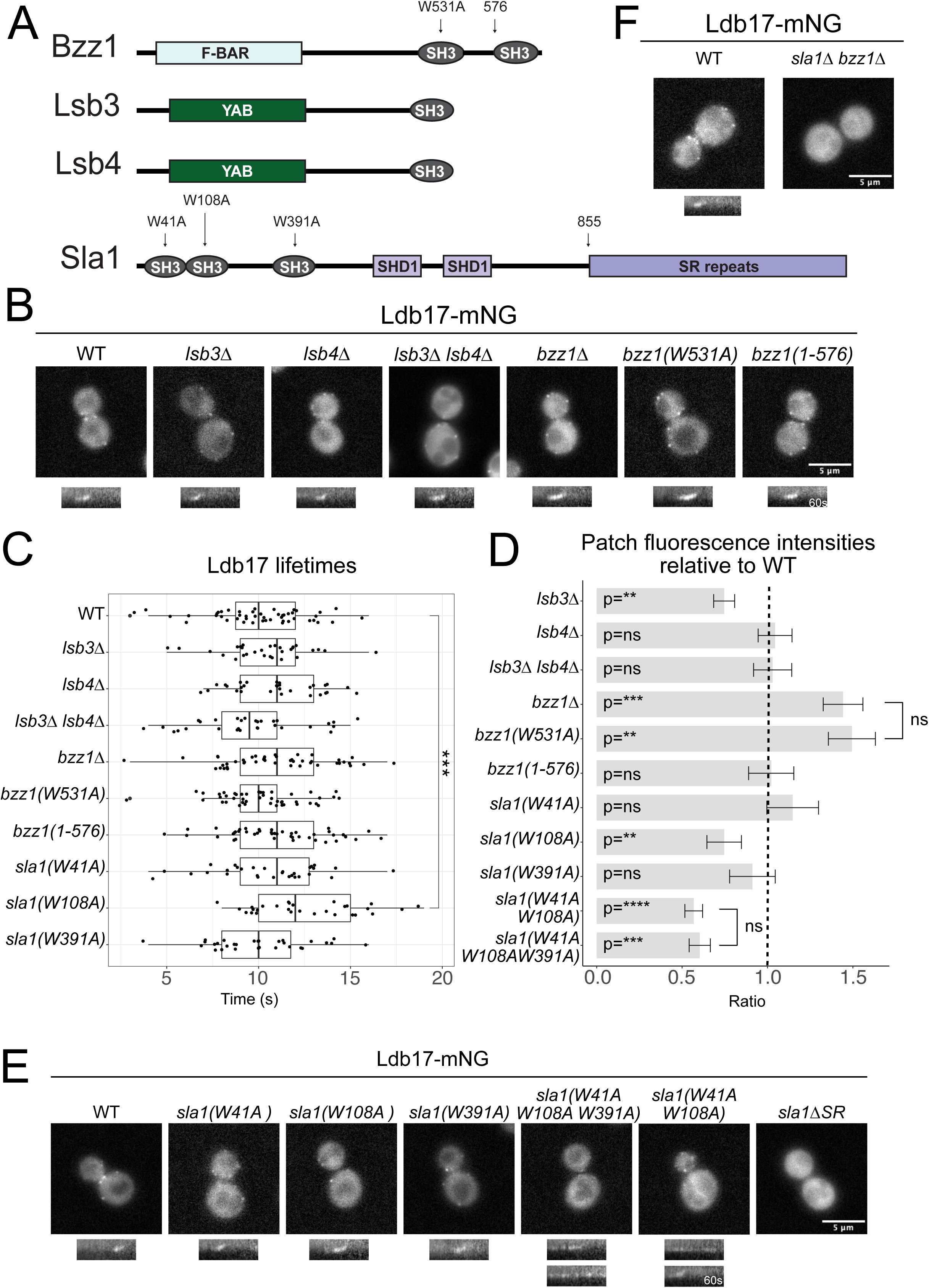
Ldb17 recruitment and regulation. (A) Structure of Bzz1, Lsb3, Lsb4 and Sla1. (B) Equatorial epifluorescence images of Ldb17-mNG in WT, *lsb3Δ*, *lsb4Δ*, *lsb3Δ lsb4Δ*, *bzz1Δ*, *bzz1(W531A)*, *bzz1(1-576)* and associated kymograph. (C) Boxplot of Ldb17-mNG lifetime in WT (n=45), *lsb3Δ* (n=30), *lsb4Δ* (n=30), *lsb3Δ lsb4Δ* (n=30), *bzz1Δ* (n=45), *bzz1(W531A)* (n=45), *bzz1(1-576)* (n=45), *sla1(W41A)* (n=30), *sla1(W108A)* (n=30) and *sla1(W391A)* (n=30). All the compared lifetimes are not significantly different except the indicated one. (D) Barplot of Ldb17-mNG patches fluorescence intensity in WT, *lsb3Δ*, *lsb4Δ, lsb3Δ lsb4Δ*, *bzz1Δ*, *bzz1(W531A)*, *bzz1(1-576)*, *sla1(W41A)*, *sla1(W108A)*, *sla1(W391A)*, *sla1(W41AW108A)* and *sla1(W41AW108AW391A)* mutants (n=30 for all). The P values on the barplot represent a U-test for the comparison between the mutants and their WT. The p values on the right of the bar plot represent a Z-test between mutants. (E) Equatorial epifluorescence images of Ldb17-mNG in *sla1(W41A)*, *sla1(W108A)*, *sla1(W391A)*, *sla1(W41A W108A)*, *sla1(W41A W108A W391A)* and *sla1(1-855)* mutants.

We designed mutants (point mutants, truncations or gene deletions) for these candidates to target specific SH3 domains (Figure 6A) and observed the consequences on Ldb17-mNG assembly dynamics. When Lsb3 and Lsb4 were deleted, either separately or in combination, it had no qualitative defect on Ldb17 recruitment to patches (Figure 6B). No major differences were also detected in Ldb17 patch lifetimes (Figure 6C). Ldb17 patch intensities in WT and *lsb4Δ* mutants were similar, but a mild decrease was detected for *lsb3Δ* mutants (Figure 6D). Bzz1 deletion resulted in Ldb17-mNG recruited to patches and internalized as WT with a similar lifetime (Figure 6B and C). However, in *bzz1Δ* cells Ldb17 patches were brighter than WT (Figure 6D), which is consistent with earlier observations (Burston et al., 2009). Bzz1 contains two SH3 domains (Figure 6A), and to identify which SH3 is behind this intensity effect, we used a point mutation in the first SH3 domain and a C-terminal truncation of the second one (Figure 6A), which were previously generated (Hummel and Kaksonen, 2023). Only the loss of the first SH3 domain resulted in brighter Ldb17 patches that matched the *bzz1Δ* null phenotype (Figure 6B, C and D). These results indicate that specifically the first SH3 domain of Bzz1 has a function that antagonizes Ldb17 recruitment to endocytic sites.

For Sla1, which contains three SH3 domains (Figure 6A), we used SH3 point mutations in the canonical PRR binding pocket, based on previous mutants (Sun et al., 2017). These SH3 domain mutants were analyzed individually or in combinations to assess Ldb17 patch dynamics. When each SH3 domain was mutated individually, *sla1(W41A)*, *sla1(W108A)* or *sla1(W391A)*, we found that Ldb17 was recruited to the membrane and internalized similar to WT in each single SH3 mutant (Figure 6E). Ldb17 intensities were unaffected in *sla1(W41A)* and *sla1(W391A)* the mutants but slightly dimmer in *sla1(W108A)* (Figure 6D). Furthermore, Ldb17 patch lifetimes of *sla1(W41A)* and *sla1(W391A)* cells did not differ from WT, but a significant increase in Ldb17 lifetime was detected in sla1*(W108A)* (Figure 6C). In combination, we found that loss of the first and second SH3 (W41A and W108A) or all three SH3 domains (W41A, W108A and W391A) domains resulted in novel Ldb17 patch behavior. In these mutants Ldb17 patches were dimmer than the WT (Figure 6D). The Ldb17 patches persisted longer than the WT at the plasma membrane, and the fluorescence signal fluctuated rapidly (Figure 6E). Due to this flickering behavior we could not determine the average lifetime of the patches. Ldb17 patches were rarely internalized in *sla1(W41A and W108A)* and never internalized in *sla1(W41A, W108A and W391A)* (Figure 6E). Together these results suggest that all three SH3 domains of Sla1 contribute to Ldb17 regulation although the second SH3 domain may have a predominant role. The Sla1 SH3 domains appear to be unessential for general Ldb17 recruitment. However, specific Ldb17 regulatory roles exist for the SH3 domains of Sla1 and the first SH3 domain of Bzz1. Ldb17 recruitment requires Sla1 (Figure 5), therefore we analyzed Ldb17 recruitment in a C-terminal deletion mutant that lacks the (Sla1 LxxQxTG repeat) SR region (Sun et al., 2017). In *sla1-ΔSR* mutants, Ldb17 patches were undetectable, similar to the null mutants (Figure 6E). Together these results suggest that the SR repeats of Sla1 are critical for recruiting Ldb17 to patches, while the SH3 domains have roles regulating Ldb17 at endocytic sites.

### Ldb17 and Dip1 differ in WASP synergy *in vitro*

We reasoned that an SH3 domain interaction with the Proline-rich tail of Ldb17 may be sufficient to activate the Arp2/3 activation activity of Ldb17. Therefore we purified the N-terminal fragment of Sla1 (1-415) and Shd1 (1-569), which contains the SH3 domains and performed fluorometric pyrene actin polymerization assays to measure the Arp2/3 activation. Addition of the SH3-Shd1 fragment to Dip1 had no detectable effect on the Arp2/3 activation activity of Dip1 (Figure 7A). Similarly, the SH3-Sla1 fragment did not affect Arp2/3 activation of Ldb17 (Figure 7A). These data suggest that the interaction between an SH3 domain of Sla1 and the proline-rich region of Ldb17 alone is not sufficient to activate Ldb17. The Sla1 SH3 domains also interact with the PRRs of Las17 (Rodal et al., 2003; Sun et al., 2017). In addition, Dip1 is proposed to synergize in *S.pombe*, which suggests a more complex relationship may exist between WASP and WDS protein, so we tested the potential combinatorial effects of Las17, WDS protein, and Sla1 ortholog. Addition of Las17 and Dip1 together resulted in a synergistic effect in actin nucleation, which was greater than either with Las17 or Dip1 alone (Figure 7B) and comparable to previous observations (Balzer et al., 2020). However, addition of the SH3 fragment of Shd1 to the two Arp2/3 activators reduced the synergistic effects (Figure 7B), possibly due to negative regulation of WASP by Sla1 (Rodal et al., 2003). Ldb17 showed no ability to synergize with WASP, and in contrast, reduced the Arp2/3 activation ability of WASP (Figure 7C). This could be due to an Arp2/3-binding competition between WASP and Ldb17. Addition of Sla1 SH3 domains to dual activator condition reduced actin nucleation similar to the *S. pombe* proteins (Figure 7C). These results suggest that there is an additional layer of regulation on the activity of *S. cerevisiae* Ldb17 compared to *S. pombe* Dip1 and that the SH3 domains of Sla1 alone are insufficient to activate Ldb17 *in vitro*.

**Figure 7:**
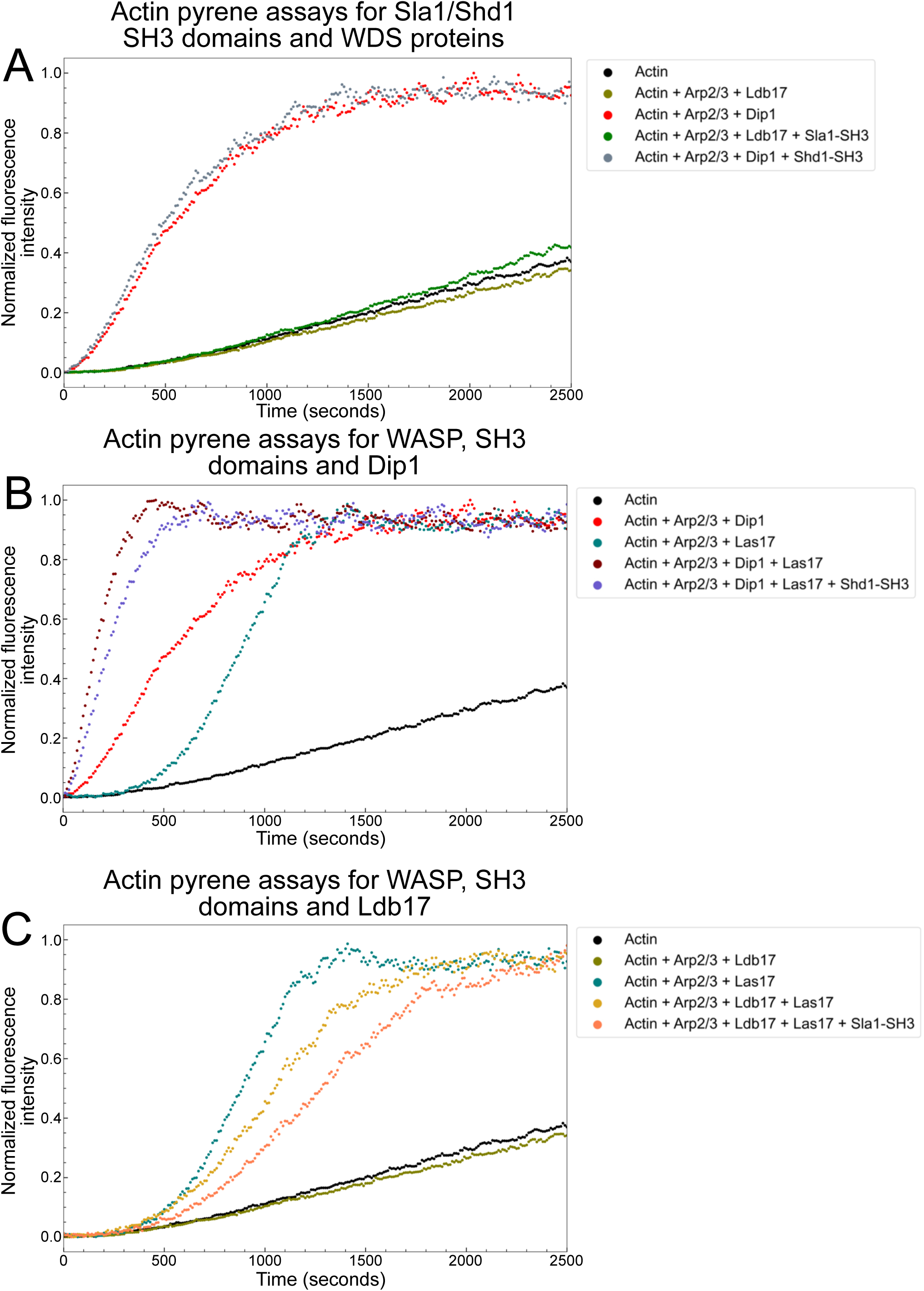
WDS protein Arp2/3 activation with WASP and Sla1 SH3 domains. (A-C) Time course of actin polymerization monitored by pyrene fluorescence. The protein concentrations are 2 µM for actin, 0.1 µM for Arp2/3 complex and 1 µM for all the other proteins. Controls are replotted from Figure 2F. Pyrene assays were performed together for fair comparison across conditions.

### The Ldb17 tail negatively regulates activity

We then wanted to investigate the function of the C-terminal tail of Ldb17 *in vivo* and its potential interaction with the SH3 domains of Sla1. We generated an Ldb17 mutant with a truncated C-terminus (a.a. 1 - 432; *ldb17-Δtail*). We then focused on the first two SH3 domains of Sla1 that have a potent effect on Ldb17 dynamics (Figure 6E). We imaged the Ldb17-Δtail mutant protein tagged with mNG in WT and *sla1*(W41A and W108A) backgrounds. We found that, similar to the full length protein, Ldb17-Δtail was recruited to patches in both conditions (Figure 8A). Quantification revealed modest reduction in Ldb17 fluorescence intensity and shortening actin patch in *ldb17-Δtail* mutant relative to WT (Figure 8C and G). Ldb17-mNG in the *sla1*(W41A and W108A) mutant was dim and flickering as previously stated and prevented determination of the beginning and end of a patch lifetime (Figure 6C and 8D). In contrast, Ldb17-Δtail-mNG in the *sla1*(W41A and W108A) mutants appeared to form stable, dim patches that internalized (Figure 8D, kymographs), but their lifetime was extended relative to WT cells (Figure 8B and D). These results are consistent with a negative regulatory role for the Ldb17 tail. The loss of the inhibitory function would not impede Ldb17 function. The Sla1 SH3 domain mutant fails to correctly regulate WT Ldb17, resulting in aberrant patch dynamics. In the double mutant, both the regulator and the regulation are removed and activity is restored. Therefore the Sla1 SH3 domains likely act via the Ldb17 tail, but the rescue is only partial, so a more complex regulation likely exists.

**Figure 8:**
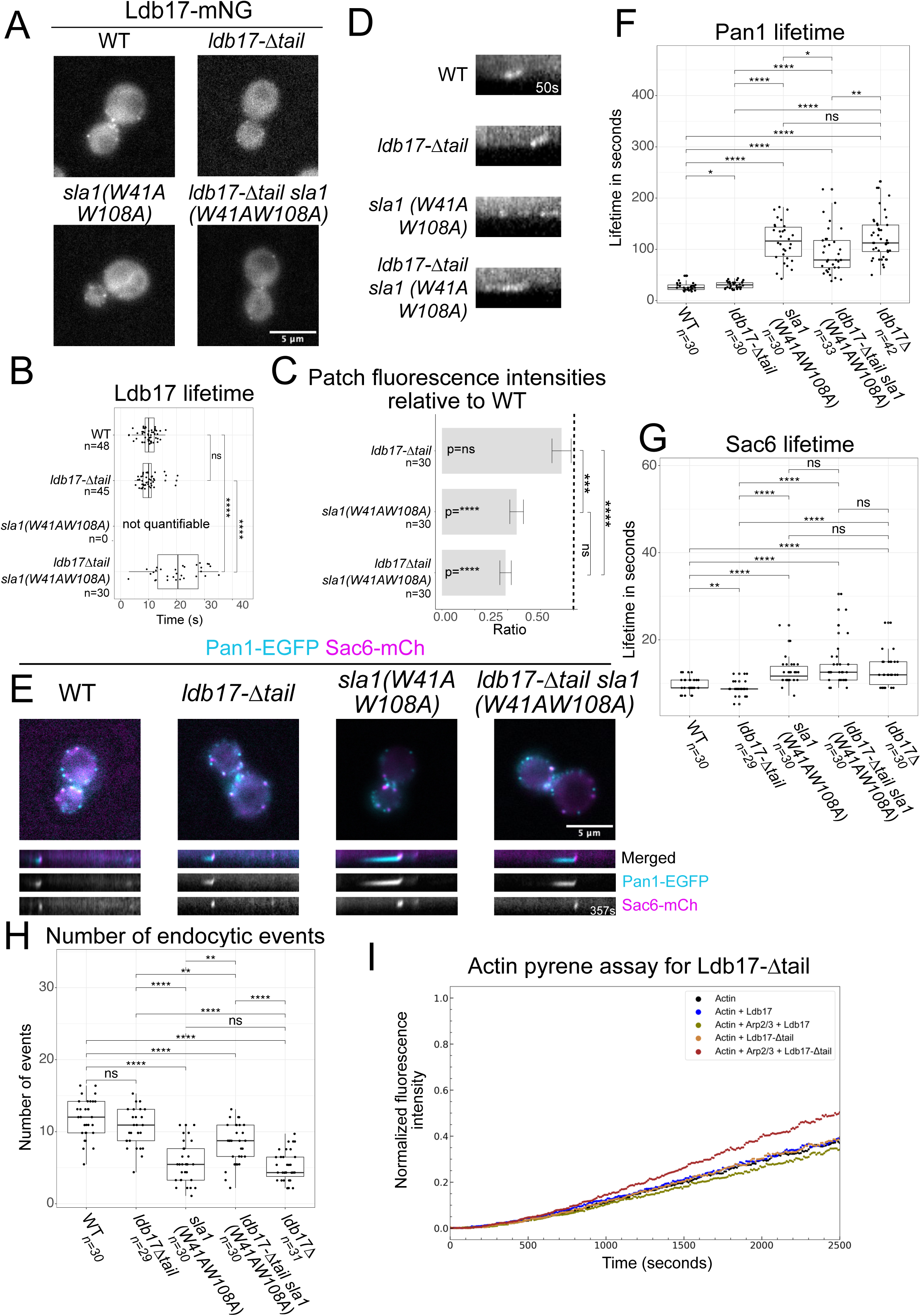
Ldb17 tail regulation. (A) Equatorial epifluorescence images of Ldb17-mNG in WT, *ldb17Δtail*, *sla1(W41A W108A)* and *ldb17Δtail sla1(W41A W108A)*. (B) Boxplot of Ldb17-mNG lifetime in WT, *ldb17Δtail*, *sla1(W41A W108A)* and l*db17Δtail sla1(W41A W108A)* cells. (C) Barplot of Ldb17-mNG patches fluorescence intensity in *ldb17Δtail*, *sla1(W41A W108A)* and *ldb17Δtail sla1(W41A W108A)* cells. The P values on the barplot represent a U-test for the comparison between the mutants and their WT. The p values on the right of the bar plot represent a Z-test between mutants. (D) Kymographs of indicated Ldb17-mNG patches. (E) *S.cerevisiae* equatorial epifluorescence images for the coat marker Pan1-EGFP, the actin marker Sac6-mCherry in WT, *ldb17Δtail*, *sla1(W41A W108A)* and *ldb17Δtail sla1(W41A W108A)* cells, and associated kymographs. (F) Boxplot of Pan1-EGFP lifetime in WT, *ldb17Δtail*, *sla1(W41A W108A)* and *ldb17Δtail sla1(W41A W108A)* cells. (G) Boxplot of Sac6-mCherry lifetime in WT, *ldb17Δtail*, *sla1(W41A W108A)* and *ldb17Δtail sla1(W41A W108A)* cells. (H) Boxplot of the number of endocytic events per cell per minute in WT, *ldb17Δtail*, *sla1(W41A W108A)* and *ldb17Δtail sla1(W41A W108A)* cells. (I) Time course of actin polymerization monitored by pyrene fluorescence. The protein concentrations are 2 µM for actin, 0.1 µM for Arp2/3 complex and 1 µM for all the other proteins. Controls are replotted from Figure 2F and Figure 7. Pyrene assays were performed together for fair comparison between conditions.

Inhibitory roles of the Ldb17 tail were further analyzed with Pan1 and Sac6 behavior in different mutants. As previously reported (Sun et al., 2017), we found that Pan1 lifetimes were extended in the *sla1*(W41A and W108A) similar to *sla1Δ* mutants (Figures 2 and 8E and F). Pan1 and Sac6 lifetimes in WT and *ldb17-Δtail* were similar (Figure 8E, F and G). In contrast the l*db17-Δtail* in combination with *sla1*(W41A and W108A) resulted in a mild suppression of the Pan1 lifetime phenotype (Figure 8E and F). A mild suppression was not detectable in the fimbrin lifetimes (Figure 8G), but the number of endocytic events was partially rescued in the double mutant (Figure 8H). Together these results further indicate that the Ldb17 tail has an inhibitory role and that the Ldb17-Δtail protein retains functionality. The partial rescue of the *sla1*(W41A and W108A) phenotype by the tailless protein supports a significant role of the Sla1 SH3 domains to relieve that inhibition. Since the rescue remains only partial, other mechanisms likely contribute to the *sla1*(W41A and W108A) mutant phenotype, possibly through an interaction with Las17 (Sun et al., 2017), or Bzz1 SH3 domain competition (Hummel and Kaksonen, 2023).

We next tested if removal of the tail could change the Arp2/3 activation ability of Ldb17 *in vitro*. A small but measurable increase in Arp2/3 activation was detected, relative to full length protein in a pyrene assay (Figure 8I), which further indicates an auto-inhibition by the C-terminal tail. The increase was dwarfed by Dip1 activity (Figure 2F and 8I). The mild *in vitro* activity further suggests that the tail may not be the only factor required to fully activate Ldb17, and aligns with our *in vivo* results. In addition, the Arp2/3 activation differences may also reflect a divergence in the actin initiation requirements of *S. cerevisiae* (pre-assembly) and *S. pombe* (co-assembly) endocytosis mechanisms.

## DISCUSSION

The WDS proteins are conserved regulators of actin assembly of opisthokonta, with the most detailed understanding of WDS protein function coming from *S. pombe* Dip1 studies (Basu and Chang, 2011; Wagner et al., 2013b; Shaaban et al., 2020; Balzer et al., 2018, 2020). These studies showed that, in contrast to WASP, Dip1 binds to and activates the Arp2/3 complex to nucleate an actin filament without the need for a pre-existing “mother filament” (Balzer et al., 2020). Recruitment of Dip1 to the endocytic site could thereby form the first actin filaments, which could then allow WASP to further activate Arp2/3 complexes and start the branched actin network assembly. Lots of work on the regulation of Arp2/3-mediated actin assembly during endocytosis has also been done in *S. cerevisiae* (Sun et al., 2006, 2017; Rodal et al., 2003). A concept of a temporally regulated actin switch was proposed for *S. cerevisiae* endocytosis (Sun et al., 2017), but the exact molecular mechanism has remained unknown. An endocytic role was demonstrated for *S. cerevisiae* Ldb17 more than 15 years ago (Burston et al., 2009), but since then the protein was largely overlooked in *S. cerevisiae* endocytosis literature, perhaps due to the low intrinsic Arp2/3 activation activity of Ldb17.

Here we show that, similar to *S. pombe*, the WDS homologs in two other fungal species (*S. cerevisiae* and *U. maydis)* localize to endocytic sites just prior to actin assembly, and that the knockout phenotype of *S. cerevisiae* Ldb17 mirrors the Dip1 deletion phenotype in *S. pombe*. These findings show that the WDS proteins share a conserved role in triggering endocytic actin assembly within fungi. Despite this functional conservation, WDS protein regulation is surprisingly different between *S. cerevisiae and S. pombe*. The Arp2/3 activation activity of Ldb17 is controlled by a mechanism that involves SH3-PRR interactions, whereas Dip1 is biochemically constitutively active. This variation of molecular mechanisms, while the cellular function is maintained, offers a basis for investigating the regulatory evolution in clathrin-mediated endocytosis.

When the actin on-switch mechanism is inactivated by mutations in WDS protein or Sla1, the timing of endocytic actin assembly appears random and cytoplasmic actin filaments may stochastically provide mother filaments for Arp2/3 activation (Basu and Chang, 2011; Chen and Pollard, 2013). Similar actin nucleation timing defects have been reported for several endocytic mutants associated with *S. cerevisiae* Sla1 and Las17 functions (*sla1Δ*, Sla1 SH3 mutants, Sla1 clathrin-binding mutants, Las17-ΔVCA mutants, Las17 phosphorylation mutants, and Pan1 mutants) (Chi et al., 2012; Feliciano and Di Pietro, 2012; Kaksonen et al., 2005; Sun et al., 2017, 2006; Tolsma et al., 2018; Tyler et al., 2021). It has been challenging to interpret these observations in a context of the biochemical functions of Sla1 and Las17 alone. Understanding the Ldb17 function in controlling actin assembly allows us to re-interpret these prior observations.

Based on our findings, we propose the following working model for the functions of *S. cerevisiae* Ldb17 in the regulation of endocytic actin assembly (Figure 9A). First, Pan1 and End3 start assembling at endocytic sites quickly followed by Las17 and Sla1 (Kaksonen et al., 2005; Sun et al., 2015). This co-arrival of proteins marks the transition from the early to the late phase of endocytosis (Carroll et al., 2012). The first two SH3 domains of Sla1 inhibit Las17 (Rodal et al., 2003). Subsequently, Bzz1, arrives at endocytic sites and its two SH3 domains compete with Sla1 for binding to a PRR of Las17 (Feliciano and Di Pietro, 2012; Hummel and Kaksonen, 2023; Soulard et al., 2005; Sun et al., 2006). Ldb17 has a unique recruitment timing among endocytic proteins just before the start of actin assembly (Burston et al., 2009; Sun et al., 2006). The recruitment of Ldb17 is dependent on the Sla1 C-terminal SR repeats, while the activity of Ldb17 is mediated by the SH3 domains of Sla1, likely via an interaction between the SH3 domains and the PRR in the Ldb17 tail. Furthermore, we found that the first SH3 domain of Bzz1 antagonizes Ldb17 assembly, possibly controlling its amount at the endocytic sites. The activation of the Arp2/3 complex by Ldb17 creates the first actin filament in close proximity to pre-assembled Las17. The transition of Sla1 SH3 domain interaction from the Las17 PRR to the Ldb17 PRR could simultaneously activate both Las17 and Ldb17. The new actin filament along with another Arp2/3 complex and Las17 initiates the assembly of branched actin network at the endocytic site to power membrane invagination (Figure 9A). The inward movement of Ldb17 with Sla1 away from Las17, could spatially separate their activities during internalization. The regulation of Ldb17 activity is likely more complex than depicted in this model, as the loss of the Ldb17 tail only partially rescued sla1 mutants and removal of the tail had mild activity relative to Dip1 on Arp2/3-mediated actin assembly (Figure 8). Furthermore, our data show that Sla1 has also Ldb17-independent activities, as the *sla1Δ* cells have unique phenotypes such as the increased coat internalization (Figure 4) and a partial co-assembly of Las17 with actin (Figure 5). Further research is needed for deciphering these mechanisms.

**Figure 9:**
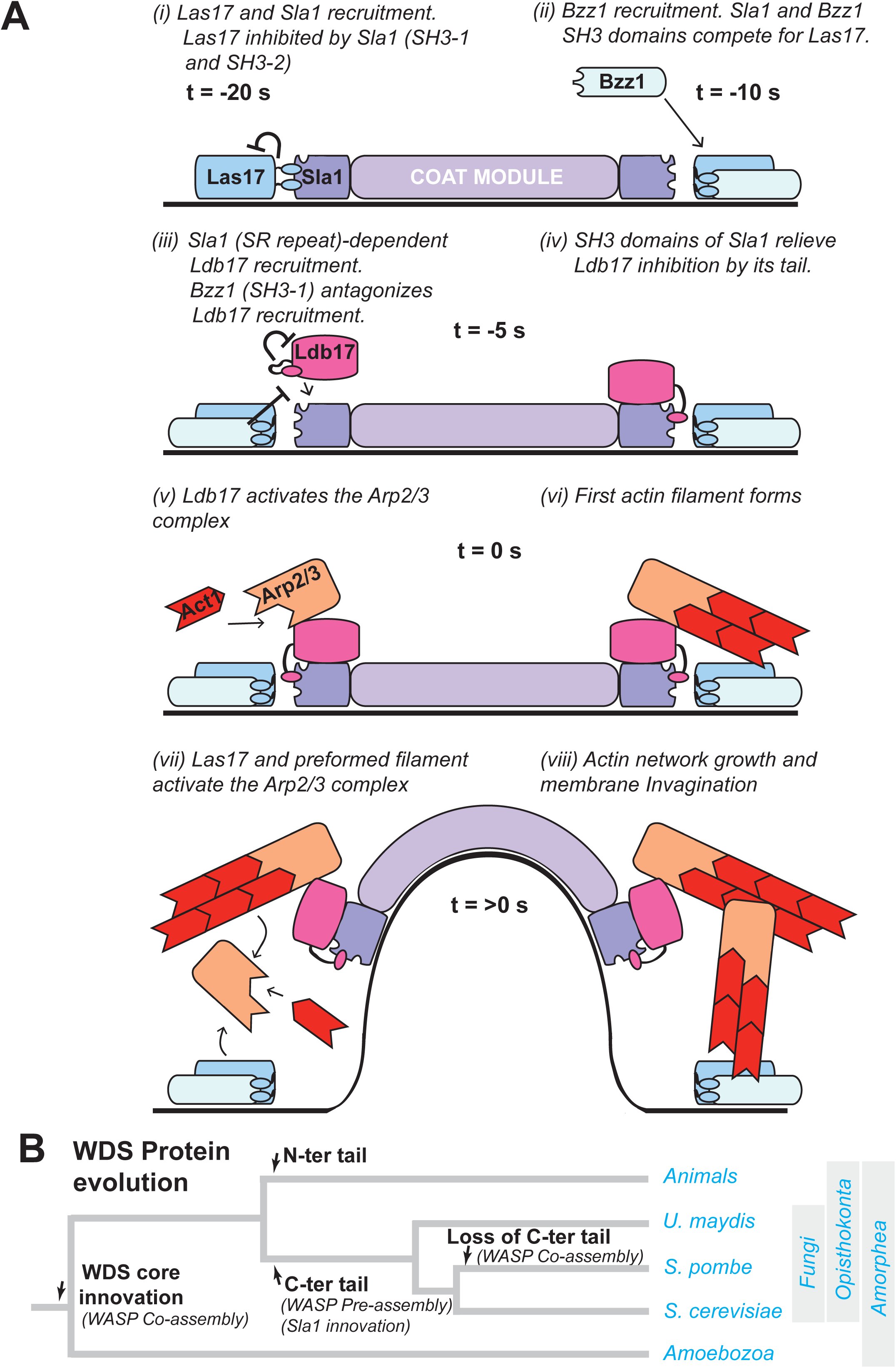
Model of Ldb17 function at budding yeast endocytic sites. (A) Schematic representation of Ldb17 recruitment and function during *S.cerevisiae* endocytosis. (B) Phylogenetic tree indicating clathrin mediated endocytic divergences.

In *S. pombe*, entry into the late phase of the endocytic pathway differs from the one depicted for *S. cerevisiae* in steps i-iv of Figure 9A. In *S. cerevisiae*, Pan1, Sla1, and WASP arrive essentially together, and then the WASP module proteins assemble in a stepwise progression (Vrp1 with Bzz1, then Myo3/5, Bbc1, and Aim21) (Kaksonen et al., 2005; Sun et al., 2006; Tonikian et al., 2009). In contrast, *S. pombe* Sla1 arrives significantly after Pan1 (Picco et al., 2024; Sirotkin et al., 2010). Sla1 is followed by the concurrent assembly of Dip1, Wsp1, Bzz1 and other WASP module-associated proteins (Sirotkin et al., 2010; MacQuarrie et al., 2019). Approximately 1s after the arrival of these proteins, Arp2/3-dependent actin assembly begins. After this point the endocytic progression in *S. pombe* is likely very similar to that of *S. cerevisiae* (Figure 9A, steps v-viii). *S. pombe* Dip1 lacks a C-terminal PRR and likely cannot interact with SH3 domains. Other differences between the Ldb17 and Dip1 structured cores may also contribute to regulatory changes (Figure 3). The differences we identified demonstrate how regulatory variation between species can inform about the mechanisms of clathrin-mediated endocytosis.

Several dynamic and mechanistic traits of clathrin-mediated endocytosis have diverged within fungal species (Picco et al., 2024; Sun et al., 2019). The WDS proteins represent another intriguing divergence. The C-terminal unstructured tail with a PRR appears in fungal WDS proteins. The appearance of the WDS tail correlates with the appearance of Sla1 early in fungal evolution (Figure 3). Deletion of the Ldb17 tail leads to subtle phenotypic changes and does not impede the actin on-switch function. This suggests that the Ldb17 tail serves a fine tuning regulatory role, but is not an integral part of the on-switch mechanism. The mild phenotype of the tail loss may have facilitated this mutation, which happened independently in a few fungal groups including *Schizosaccharomyces* (Figure 3). Both *U. maydis* and *S. cerevisiae* pre-assemble WASP prior to actin assembly at endocytic sites and have a C-terminal tail in their WDS proteins, while *S. pombe* co-assembles a WASP together with actin and has a tailless Dip1 (Figure 9B). It is an intriguing possibility that the WASP assembly mode and WDS protein tail status are causally linked. The loss of the regulatory tail could have facilitated the evolution of the less elaborate, simultaneous assembly of WASP and its regulators together with actin. Alternatively, the loss of the WDS tail became possible with the changes in WASP assembly. These changes may allow *S. pombe to* generate large actin patches and longer coat movement relative to other species (MacQuarrie et al., 2019; Sun et al., 2019; Picco et al., 2024). Interestingly, the holozoan Spin90 has an analogous unstructured tail N-terminally attached to the conserved WDS core, which has both PRRs and a SH3 domain. This suggests that the PRR-SH3-based regulation may have been invented independently in animals and fungi. Shared features between endocytic pathways like, WASP co-assemby (Mammalian cells and *S. pombe*) and SH3-PRR regulations (Mammalian cells and *S. cerevisiae*) offer opportunities to dissect how clathrin-mediated endocytosis evolves mechanistically, but still preserve process functionality.

## MATERIALS AND METHODS

### Strains and plasmids

All strains in this study are listed in Table S1. The strains were generated either by transformation using homologous recombination or by mating, sporulation and subsequent tetrad dissection. General yeast growth, transformation and mating manipulations were done according to (Dunham et al.; Hagan et al., 2016). For the *S.cerevisiae* transformation, constructs were made by PCR amplification of a DNA fragment containing a selection cassette (Janke et al., 2004), and then confirmed by colony PCR and sequencing. Internal point mutations of Sla1 were generated by first constructing the desired mutant gene in a pMK0045 plasmid using PCR mutagenesis. The mutated Sla1 sequence was then amplified by PCR using long overlapping primers. For *U. maydis* and *S. pombe* transformations plasmid constructs were built with up to 1000 bp flanking regions, linearized and integrated into the desired locus and verified by PCR as described above (Bösch et al., 2016). All plasmids used in this study are listed in Table S2.

### Media, buffers and growth conditions

All the media used are kindly provided by the biochemistry department (Brigitte Bernadets) or by Anne-Sophie Rivier-Cordey for the Kaksonen group. They were prepared following the standard protocols (Treco and Lundblad, 1993). All media formulations are listed in Table S3. *S. cerevisiae* strains were grown at 24 or 30 °C on 2% agar plates supplemented with YPD. *S. pombe* and *Ustilago maydis* strains were grown at 24°C on 2% agar plates supplemented with YES and YEPS-light respectively

### Protein purification

#### Actin purification and labeling

Actin was isolated from rabbit muscle acetone powder (Pel-Freez Biologicals) (Spudich and Watt, 1971), gel-filtered on Superdex 200 column and stored as monomers at 4°C in G-buffer (5 mM Tris-HCl, ph 8.0; 0.2 mM ATP; 0.1 mM CaCl_2_; 0.5 mM DTT). Actin monomers were labeled at Cys-374 with pyrene-iodoacetamide (Kouyama and Mihashi, 1981). Pyrene-labeled actin monomers were mixed with unlabeled actin monomers to achieve 5% labeling.

#### Arp2/3 complex purification

*S. cerevisiae* Arp2/3 complex was purified from commercially purchased baker’s yeast (Kastalia, LESAFFRE France) as described in (Antkowiak et al., 2019).

#### Las17 purification

S. cerevisiae Las17 (375-Cter), containing the PRR and C-terminal WCA domain, was expressed and purified from Rosetta 2(DE3)pLysS cells as described in (Boiero Sanders et al., 2022).

#### Ldb17 and variants purification

The genes of Ldb17 and its variants were amplified from the *S. cerevisiae*’s genome and cloned into pCoofy6 vector (A gift from Sabine Suppmann, Addgene plasmid #43990) using a ligation-independent cloning approach (Scholz et al., 2013). Proteins were expressed in Rosetta 2(DE3)pLysS cells in Autoinduction LB media (Formedium #AIMLB0210) overnight in +20°C. Cells were lysed by sonication in lysis/wash buffer (20 mM Tris-HCl pH 8.0; 140 mM NaCl; 1 mM DTT; 40 mM Imidazole; 1mM PMSF; cOmplete EDTA-free protease inhibitor tablets from Roche) 6 times 20 seconds on ice. Lysates were clarified by centrifugation and soluble fractions were filtered through 0.45 µm filter and loaded on a 5 mL HisTrap HP column (Cytiva #17524802) using an Äkta pure system (Cytiva). The affinity column was washed with the lysis/wash buffer and eluted with an elution buffer (20 mM Tris-HCl pH 8.0; 140 mM NaCl; 1 mM DTT; 250 mM Imidazole). Peaked fractions were pooled and 30 µg/ml of Senp2 protease was added. The solution was dialyzed overnight at 4°C against dialysis/gel filtration buffer (20 mM Tris-HCl pH 8.0; 100 mM NaCl; 1 mM DTT), filtered through a 0.22 µm filter and concentrated to 500µl-2ml in an Amicon-Ultra concentrator. The sample was loaded onto a Superdex 200 Increase 10/300 GL column (Cytiva #28990944), previously equilibrated with the dialysis/gel filtration buffer. Peak fractions were pooled together, concentrated in an Amicon-ultra concentrator, flash-frozen in liquid nitrogen and stored at -80°C.

#### Shd1 and Sla1 SH3 domains purification

Sla1 and Shd1 SH3 domains DNA sequences with an N-terminal SUMO3 tag were synthesized at GenScript and integrated into pET28a(+) vector for recombinant protein expression. A protocol similar to that used for the purification of Ldb17 variants was followed for the affinity purification, with the exception that the lysis/wash buffer contained 20 mM Hepes, pH 7.5; 150 mM NaCl; 1mM DTT; 30mM imidazole; 1 mM PMSF and cOmplete EDTA-free protease inhibitor tablets from Roche, that the elution buffer contained 20 mM Hepes, pH 7.5; 150 mM NaCl; 1 mM DTT; 300 mM imidazole, and that the dialysis/gel filtration buffer contained 20 mM Hepes, pH 7.5; 150 mM NaCl; 1 mM DTT. The purified proteins were dialysed overnight against dialysis buffer 2 (20 mM Hepes pH 7.5; 75 mM NaCl; 1 mM DTT) and loaded onto a HiTrap Q HP anion exchange column (Cytiva #17115401). The proteins were eluted with the ion exchange elution buffer (20 mM Hepes pH 7.5, 500 mM NaCl, 1 mM DTT) and peak fractions were flash-frozen in liquid nitrogen and stored at -80°C.

### Pyrene-actin polymerization assays

Pyrene-labeled Ca-ATP-actin monomers were incubated for 5 minutes on ice in the presence of 0.2 mM EGTA and an 11-fold molar excess of MgCl_2_ over actin. Actin polymerization was induced by the addition of 1/10 volume of 10x KMEI (1x contains 50 mM KCl; 1 mM MgCl2; 1 mM EGTA; and 10 mM imidazole-HCl, pH 7.0). Changes in pyrene fluorescence over time were recorded on a Safas Xenius XC spectrofluorimeter with an excitation at 365 nm and a detection at 407 nm. All experiments were reproduced at least 3 times. The curves presented in panels 2F, 7A, 7B, 7C and 8I come from the same experimental data set.

### Phylogenetic, sequence alignment analysis

Orthologies were mined with PSI-BLAST and blastp at https://www.ncbi.nlm.nih.gov/. WDS core and tail regions were defined using MUSCLE and COBALT alignment tools. Proline rich regions were identified by sequence gazing tail regions. Structures were visualized in PyMOL.

### Microscopy

#### Sample preparation

For *Saccharomyces cerevisiae,* strains were grown overnight at 24°C in 5 mL synthetic complete medium without l-tryptophan (SC-TRP) culture in a shaker. Then, they were diluted to OD_600_ = 0.2 and grown until OD_600_ reached between 0.4 and 0.8. Ahead of time, coverslips were prepared by placing 50µL of Concanavalin IV (ConA, Sigma-Aldrich, 4 mg/mL in H2O) on the coverslip for 20 min, washing three times with 40 µL SC-trp and dropping 40 µL of cell culture on the ConA coated coverslip for 20 min. Then, the cell drop was washed three times and a final 40 µL drop of Sc-trp was put on the surface. For drug-treated cell imaging, the cells were washed and then incubated with 40 µL SC-trp media containing 0.2 mM LatA (Enzo), or CK-666 (Sigma-Aldrich) or DMSO (Sigma-Aldrich) solution for 15 min. The cells were imaged within 30 min after drug treatment.

In Schizosaccharomyces *pombe* and *Ustilago maydis,* strains were grown overnight at 24°C in 10 mL EMM in a shaker. The day after, they were diluted to OD_600_=0.02 in 10 mL EMM for Schizosaccharomyces *pombe* and 0.01 for *Ustilago maydis* and put back overnight at 24°C while shaking. The next day, strains were imaged at OD_600_ = 0.4-0.8. Then, 200 µL of heated agarose with 2%EMM is pressed between two glass microscope slides separated by two thin spacers about 0.5 mm thick. While drying, 200 µL of cell culture is spun 1 min at 3000 rpm and resuspended in 5-7 µL of the culture. Then, a 1 µL drop is put on the EMM 2% agarose, a coverslip is added and sealed by pouring Valap on the edges. For drug-treated cell imaging, the 5-7 µL pelleted cells from a log phase culture were incubated with 40µL EMM containing 0.2 mM LatA, or CK-666 or DMSO solution for 15 min before being pipetted on the 2% EMM agarose pad that contained the corresponding drug concentration to avoid diffusion. The cells were imaged within 30 min after this incubation period.

#### Data acquisition

Widefield images were acquired with an Olympus IX81 widefield microscope equipped with a 100×/NA1.45 objective and an ORCA-ER CCD camera (Hamamatsu). The illumination source used was an X-CITE 120 PC (EXFO) metal halide lamp. The excitation and emission light when imaging EGFP-, mNeonGreen- and mCherry-tagged proteins were filtered through the U-MGFPHQ and U-MRFPHQ filter sets (Olympus). The microscope was controlled using Micromanager (Edelstein et al., 2014).

TIRF images were acquired on an Olympus IX83 widefield microscope equipped with a 150×/NA1.45 objective and an ImageEM X2 EM-CCD camera (Hamamatsu) and controlled by the VisiView software (Visitron Systems). A 488 nm and 561 nm lasers were used for the excitation of EGFP-, mNeonGreen- or mCherry-tagged protein,. Excitation and emission were filtered using a TRF89902 405/488/561/647 nm quad-band filter set (Chroma). Laser angles were controlled by iLas2 (Roper Scientific)

#### Data analysis

General image analysis was performed using the Fiji distribution of ImageJ (Schindelin et al., 2012; Rueden et al., 2017). All display images were corrected for background fluorescence using the rolling ball algorithm of ImageJ, and movies were corrected for photobleaching using a custom ImageJ macro. For particle detection and tracking, the Particle Tracker (Sbalzarini and Koumoutsakos, 2005) of the MosaicSuite was used. Average trajectories were obtained as described previously (Picco *et al*., 2015). Plots and statistical analyses were computed using R. The statistical test used is the Wilcoxon-Mann-Whitney for the pairwise comparison: ns for P-value > 0.05, * for P-value ≤ 0.05, ** for P-value ≤ 0.01, *** for P-value ≤ 0.001 and **** for P-value ≤ 0.0001.

Patch lifetimes: Lifetimes were measured from dividing yeasts and computed for each strain from at least three different fields of view. The number of frames from the appearance to the disappearance of a patches was counted and multiplied by the time interval between frames measured in the metadata.

Number of endocytic events: we counted the number of sites showing the colocalization of a coat marker with an actin marker followed by their internalization. This sequence of events illustrates a successful endocytic event.

Tracking and fluorescence intensity profile: A method based on the averaging of subpixel centroid positions was used (Picco and Kaksonen, 2017). For each protein tracked, the centroid trajectory is recorded for multiple endocytic events and at least 35 trajectories are averaged.

This method also provides the fluorescence intensity over time. Fluorescent intensity profiles of Las17-EGFP are aligned using the alignment of the peak of Sac6-mCherry using a python script (Picco et al., 2024).

Patch fluorescence intensity: Cell cytoplasm was subtracted from the original image using the Fiji median filter. Fluorescence intensity of patches from at least 3 fields of view was recorded using the circle tool of Fiji with a 10 pixel radius. Fluorescence intensity from the background around the patch was also recorded. After averaging the patch fluorescence intensity value and the background fluorescence intensity value, the background value was used to normalize the patch value.

The plots were rendered using the ggplot package of R or Python 3.11.

## Acknowledgements

We thank Daniel Hummel, Anne-Laure Boinet, Mamta, Yushi Jiang, and Bagyashree V. T. for critical reading of the manuscript and helpful discussions. This work was supported by the Swiss National Science Foundation grant to M.K. (310030_212288).

## SUPPLEMENTAL TABLES

**Table S1:**
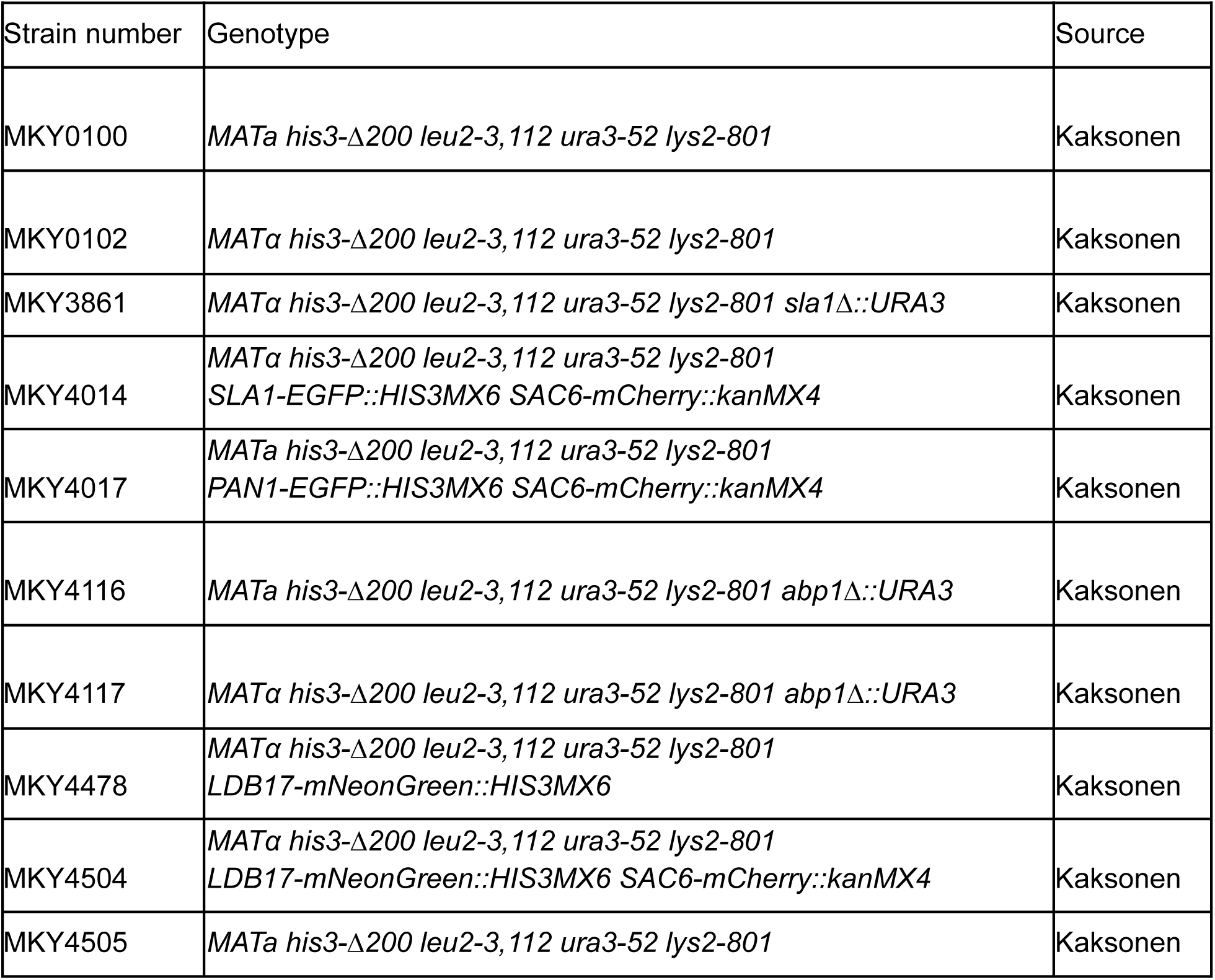
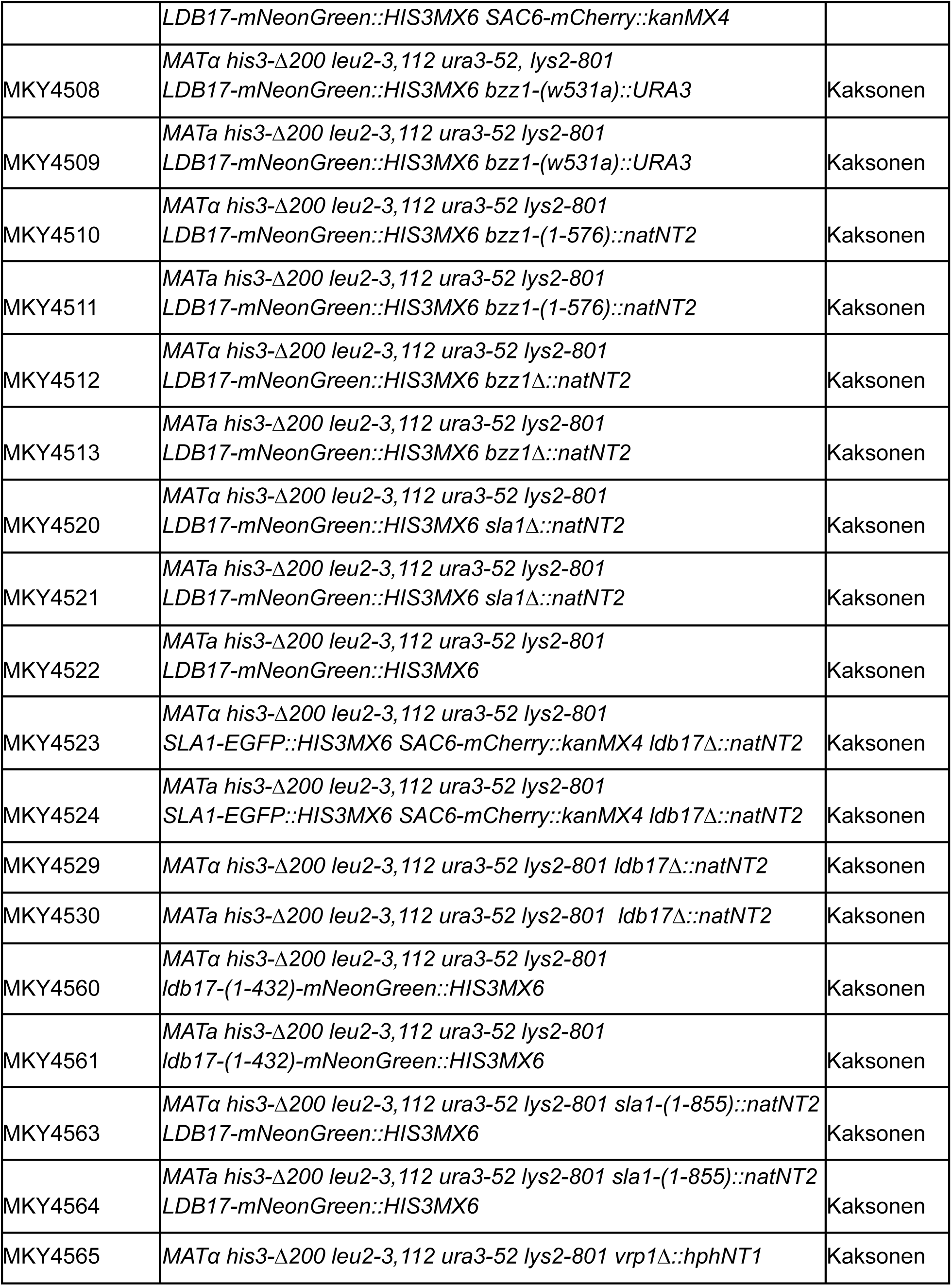
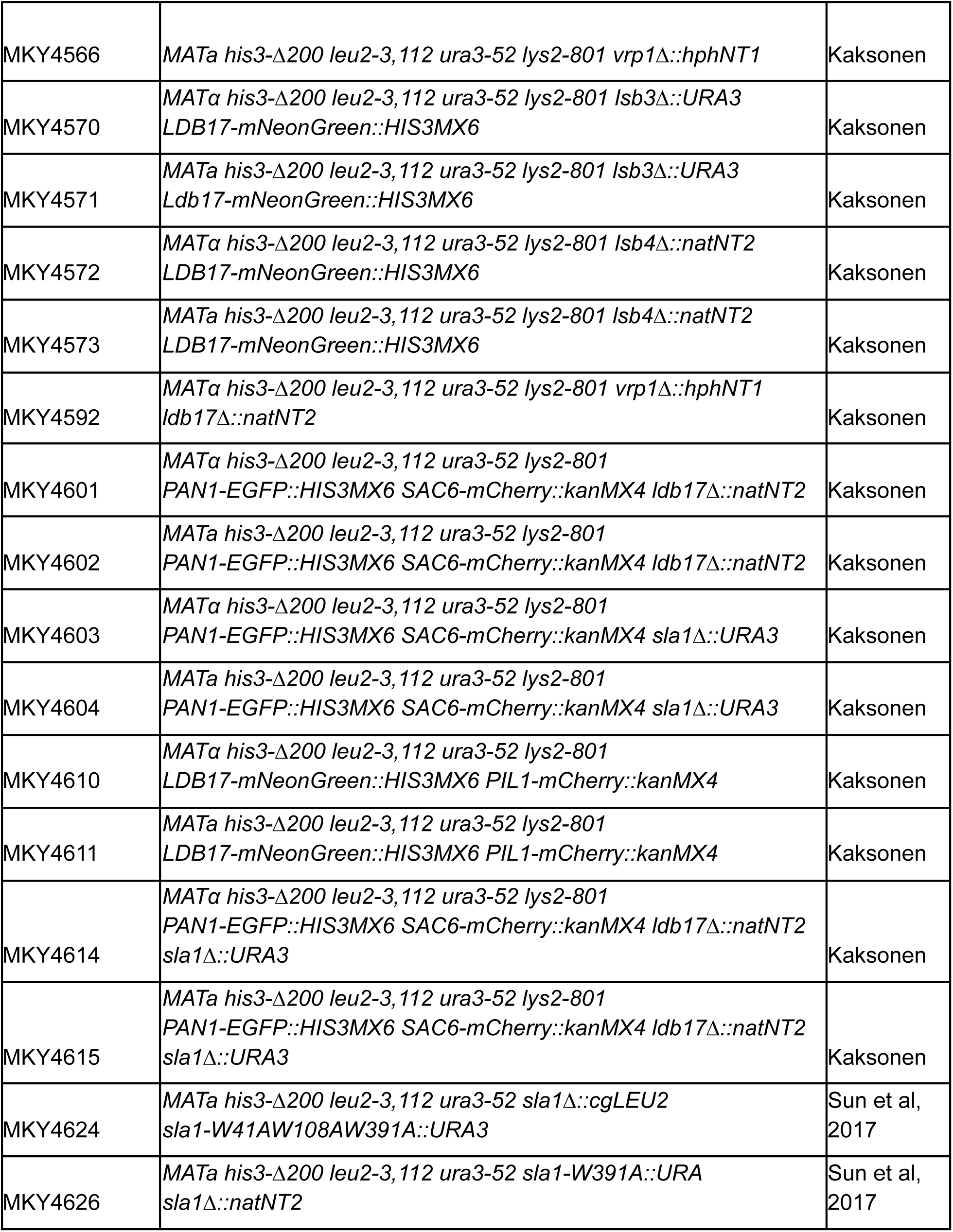
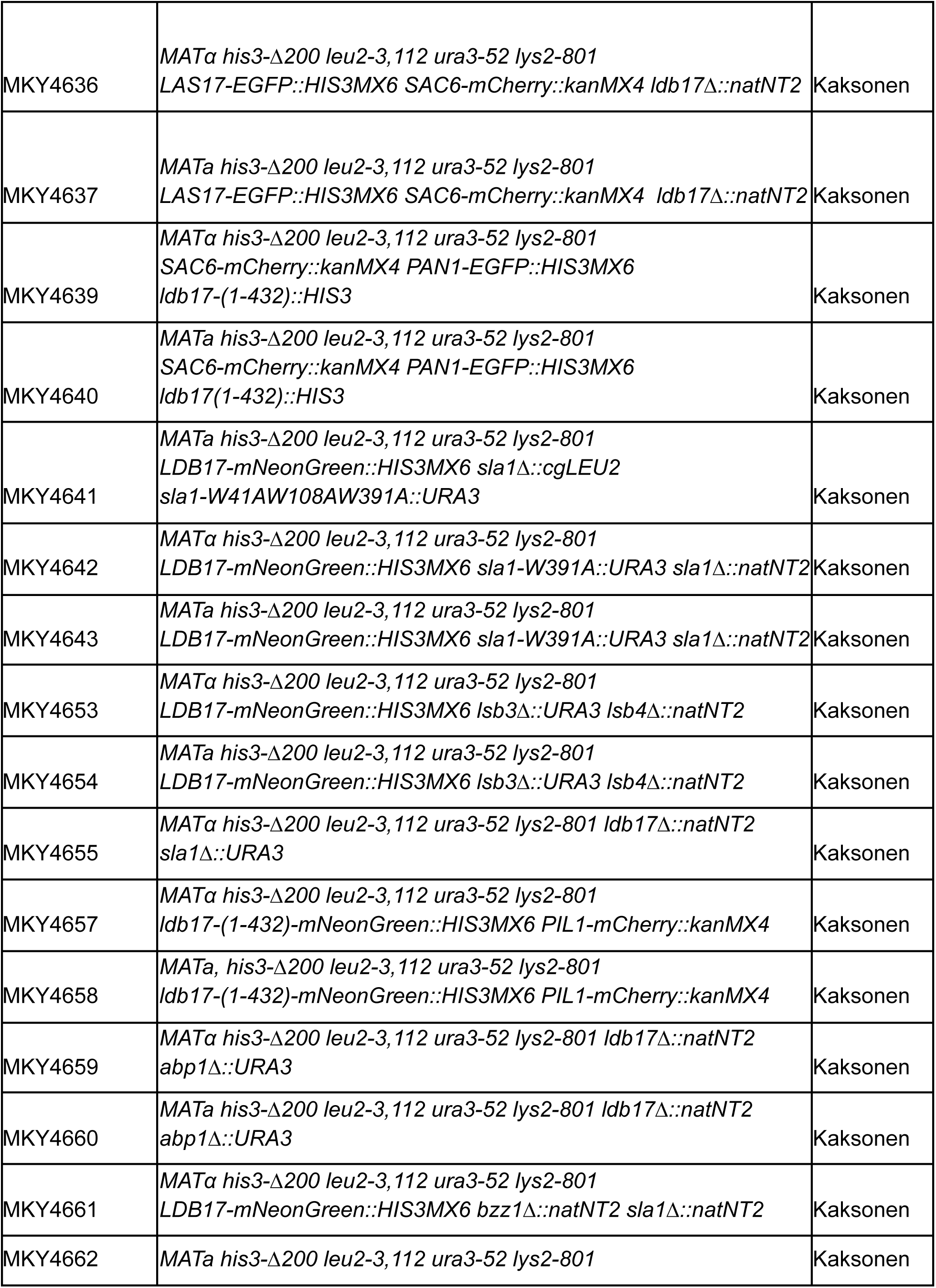
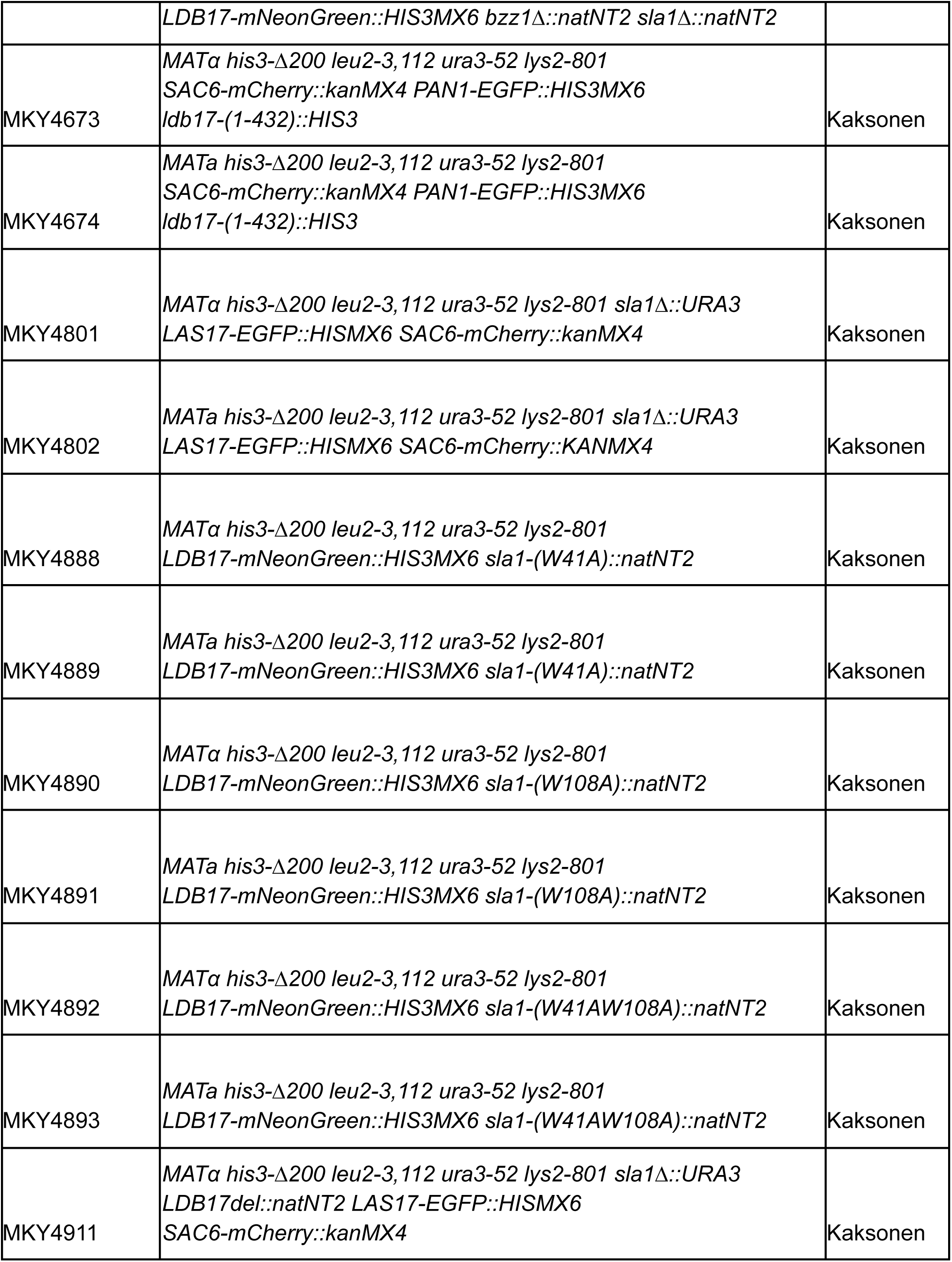
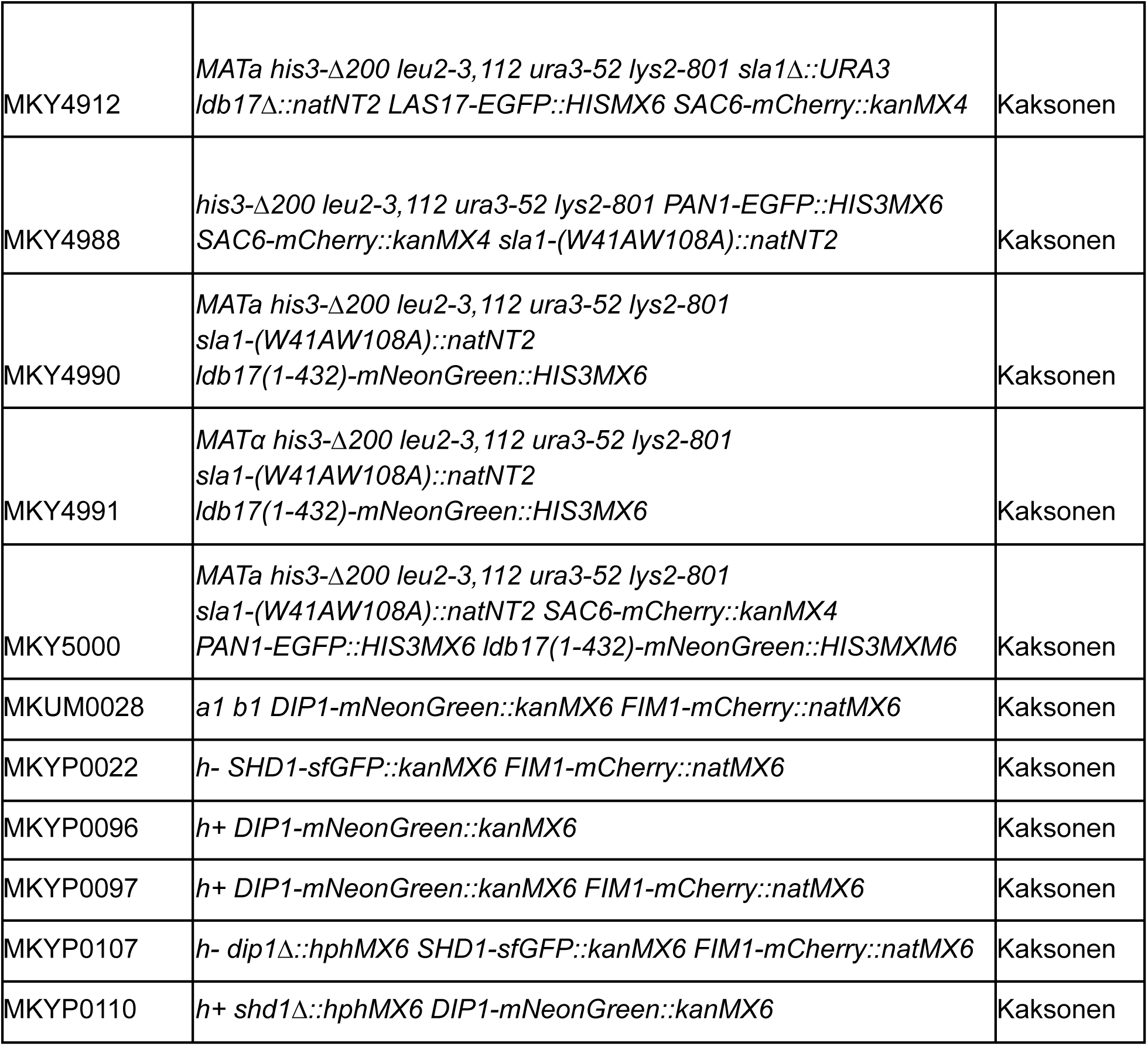
Yeast strains.

**Table S2:**
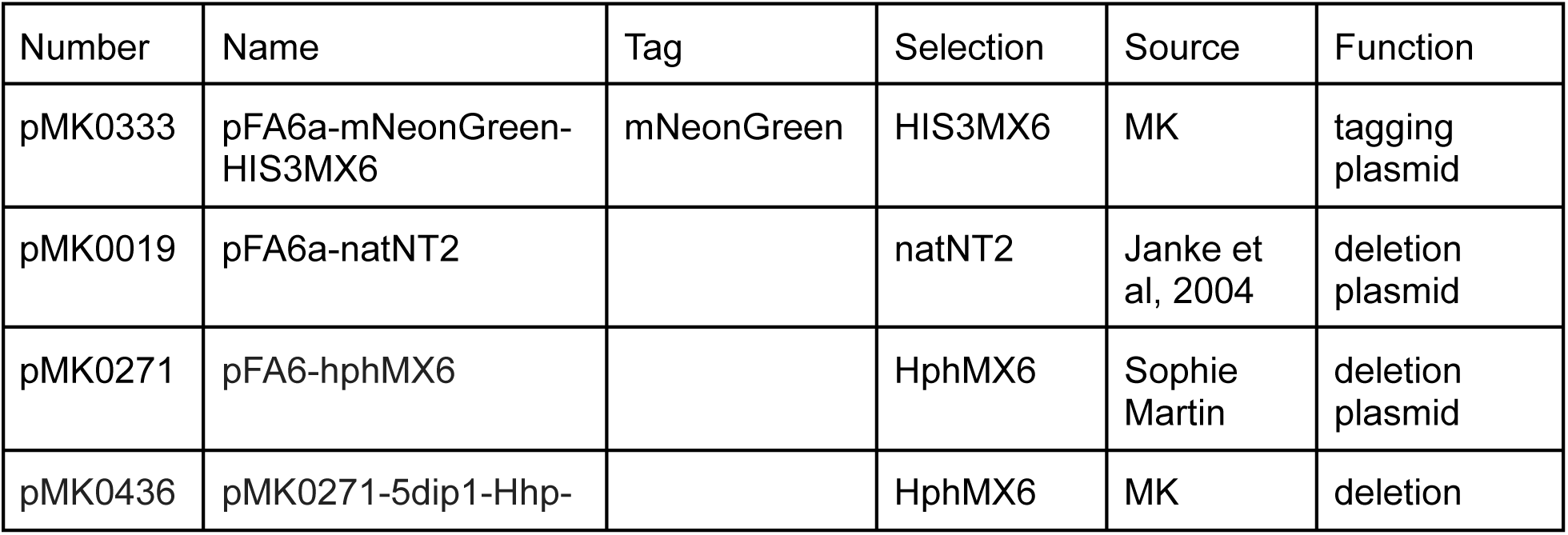
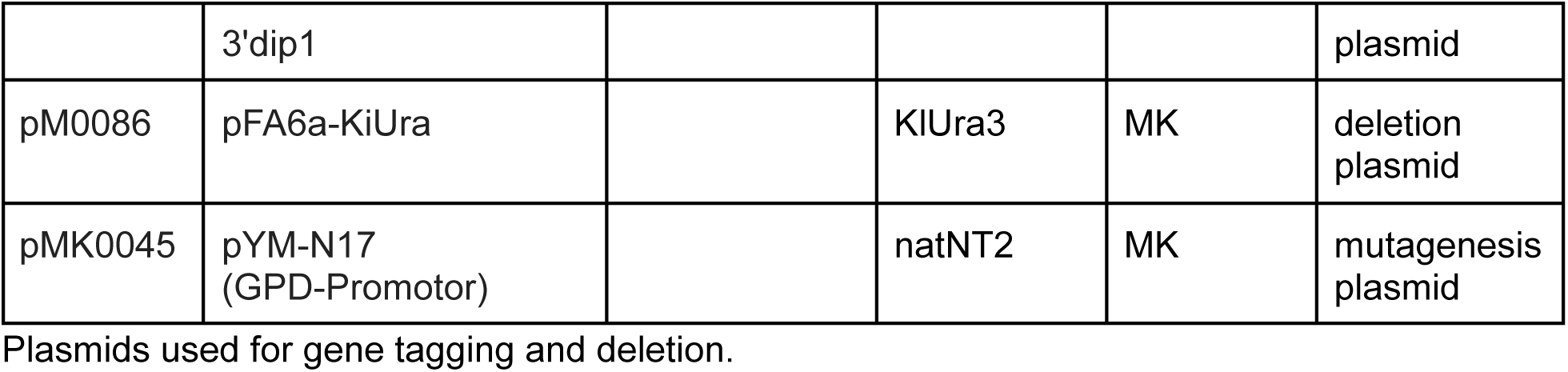
Plasmids.

**Table S3:** Yeast media and buffers.

| Media | Species | Components (w/v) |
| --- | --- | --- |
| YPD | *S. cerevisiae* | 1% Yeast extract, 2% Bacto peptone, 2% Glucose, 40 mg/L uracil & adenin |
| YES | *S. pombe* | 0.5% Yeast extract, 3% Glucose, 0.0225% Adenine, 0.0225% Histidine, 0.0225% Leucine, 0.0225% Lysine, 0.0225% Uracil |
| YEPS-light | *U. maydis* |  |
| Sc-Trp | *S. cerevisiae* | 0.2% Sc-Trp, 0.67% yeast nitrogen base, 2% glucose |
| EMM | *S. pombe*<br>*U. maydis* | 0.3% potassium phthalate monobasic, 0.22% sodium phosphate dibasic, 0.5% ammonium chloride, 2% anhydrous glucose, 2% salts, 1% vitamins, 0.1% minerals |
| PEG | *S. cerevisiae* | 100nM LiAc, 10 mM Tris-HCl pH8, 1 mM EDTa pH8, 10% PEG |
| SORB | *S. cerevisiae* | 100nM LiAc, 10 mM Tris-HCl pH8, 1 mM EDTa pH8, 1 M Sorbitol |

Medias used in this study. For plates 2% agar was added to these media.

## SUPPLEMENTAL FiGURES AND TEXT

**Supplemental Figure S1:**
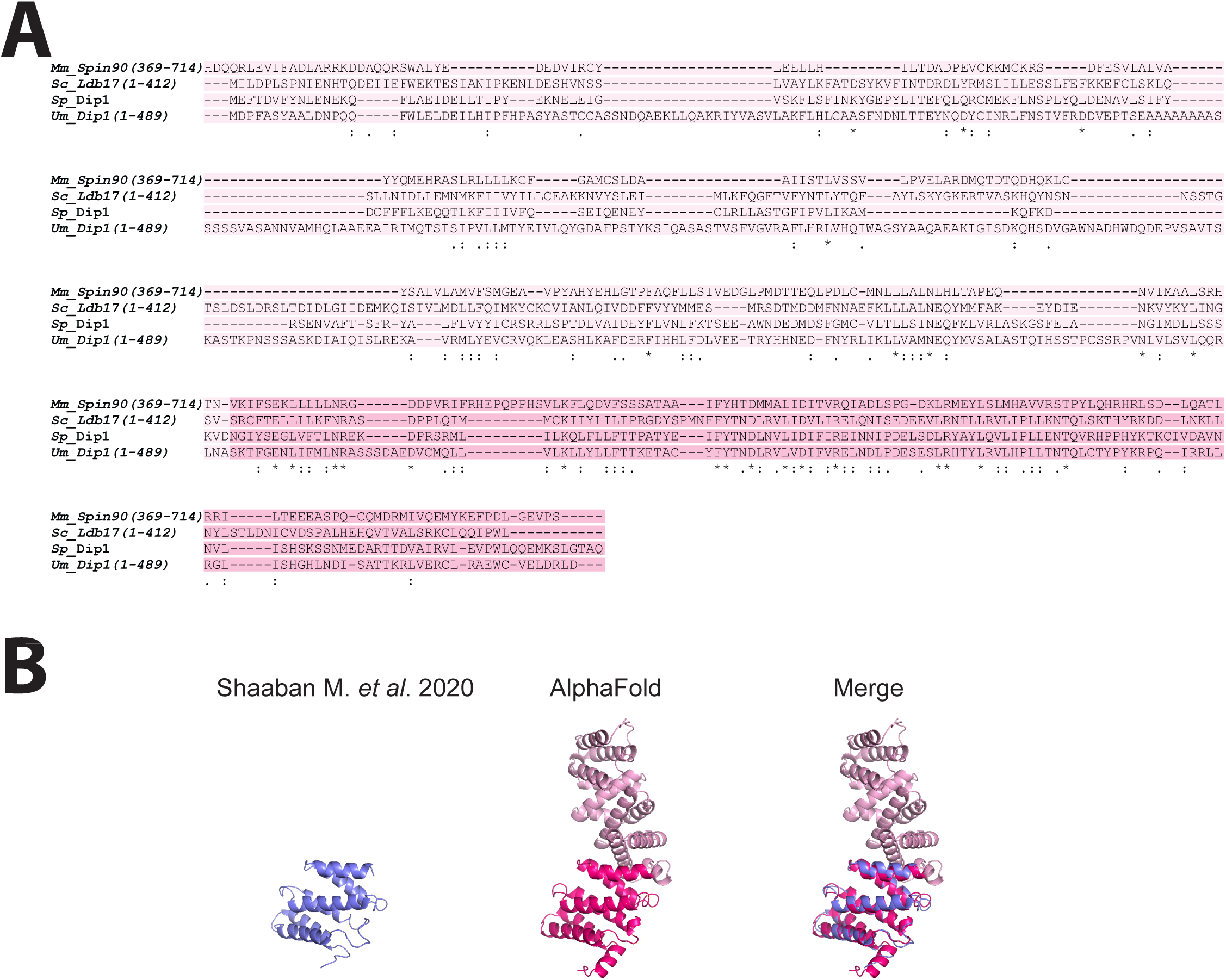
(A) *S. pombe* Dip1 structure (https://www.ncbi.nlm.nih.gov/pmc/articles/PMC8717925/) and Alpha fold prediction for *S. pombe* Dip1. Variable region of the WDS core regions (light blue) and constrained region (dark blue). Both structures were aligned and oriented with PyMOL. (B) MUSCLE alignment of indicated WDS orthologs regions. WDS structured core beginning and end were identified using Alphafold with variable region (light blue) and constrained region (dark blue).

**Supplemental Text:** Text file that lists WDS protein accession numbers and sequences, with features highlighted in indicated colors.

## SUPPLEMENTAL TEXT

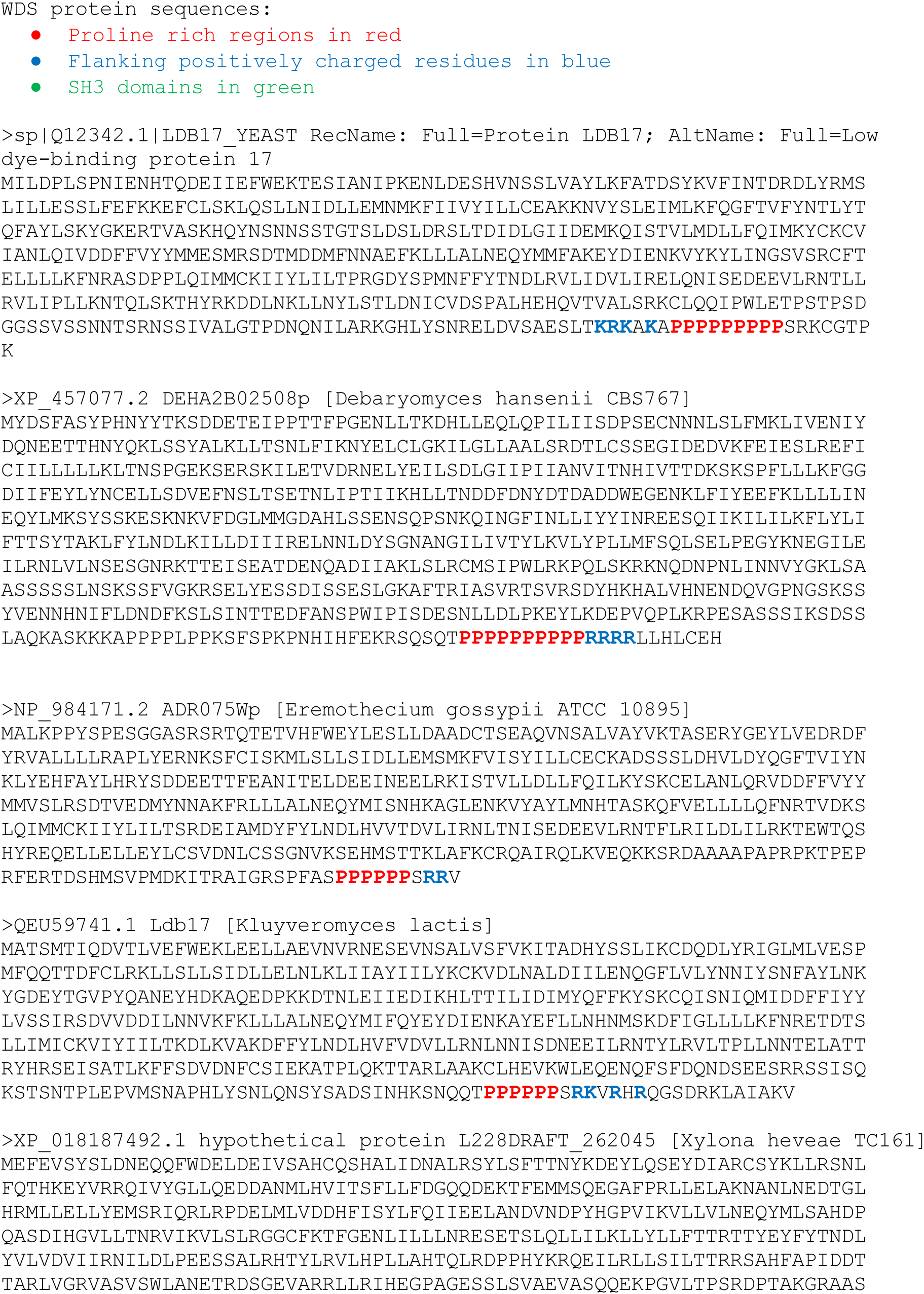
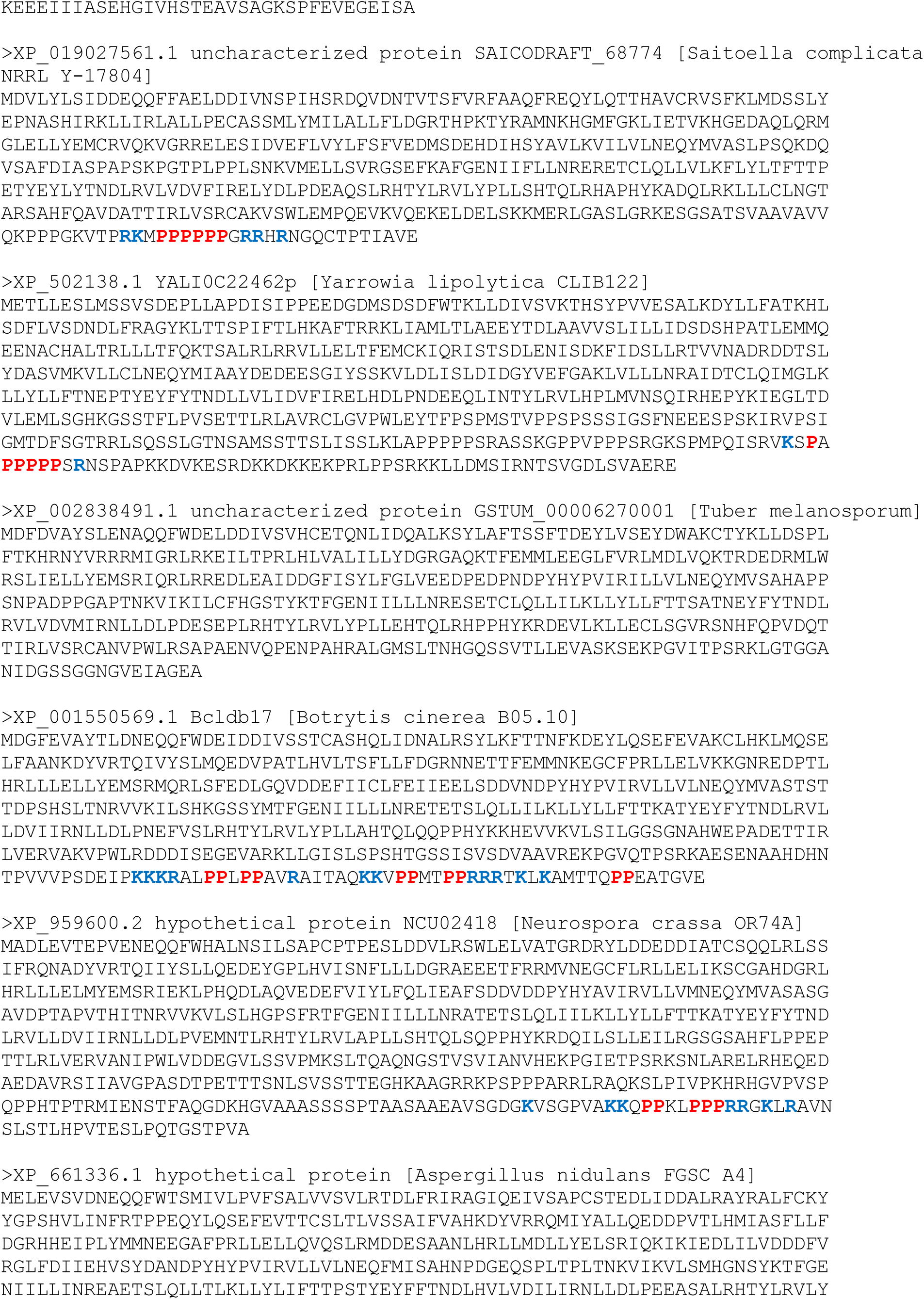
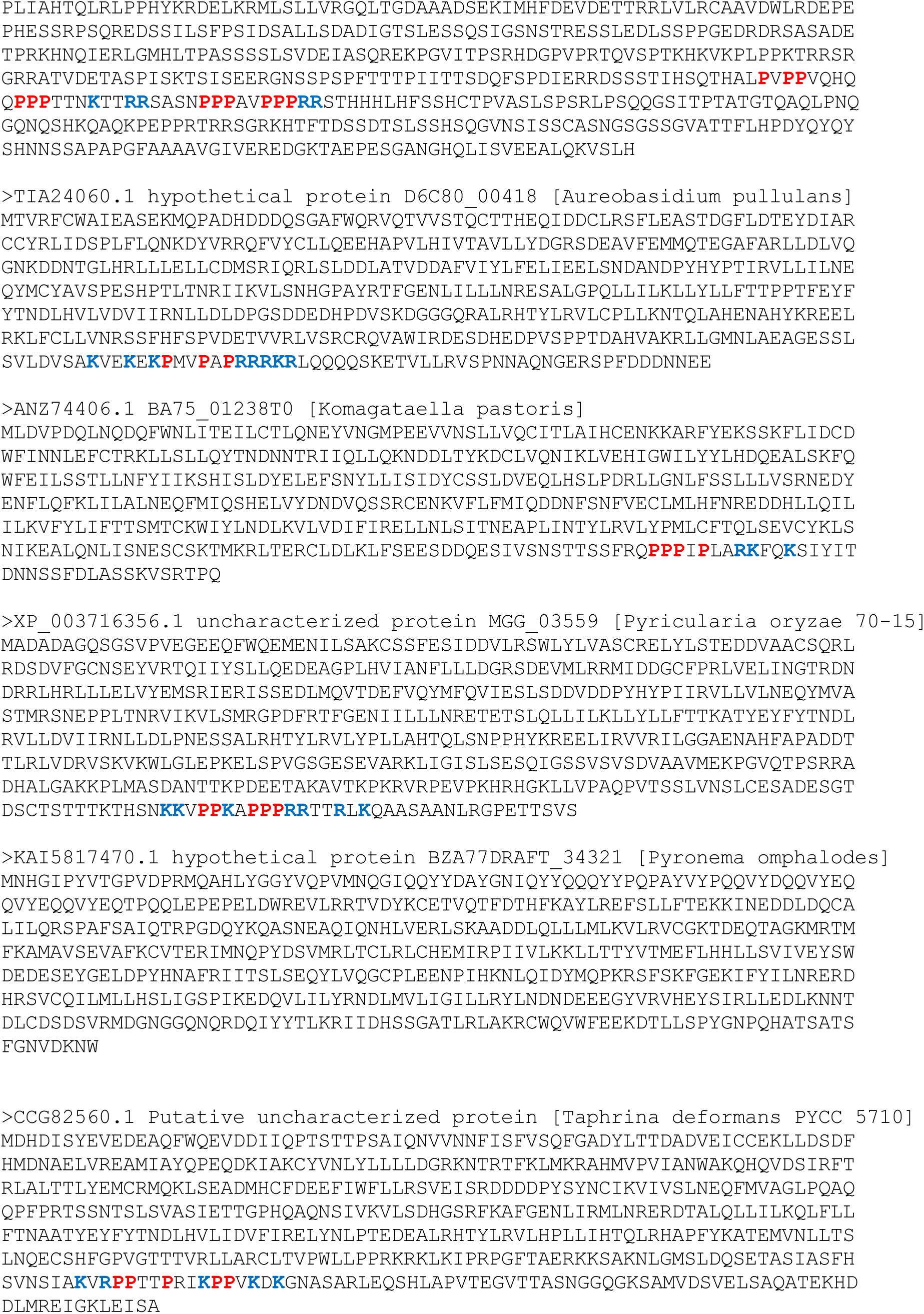
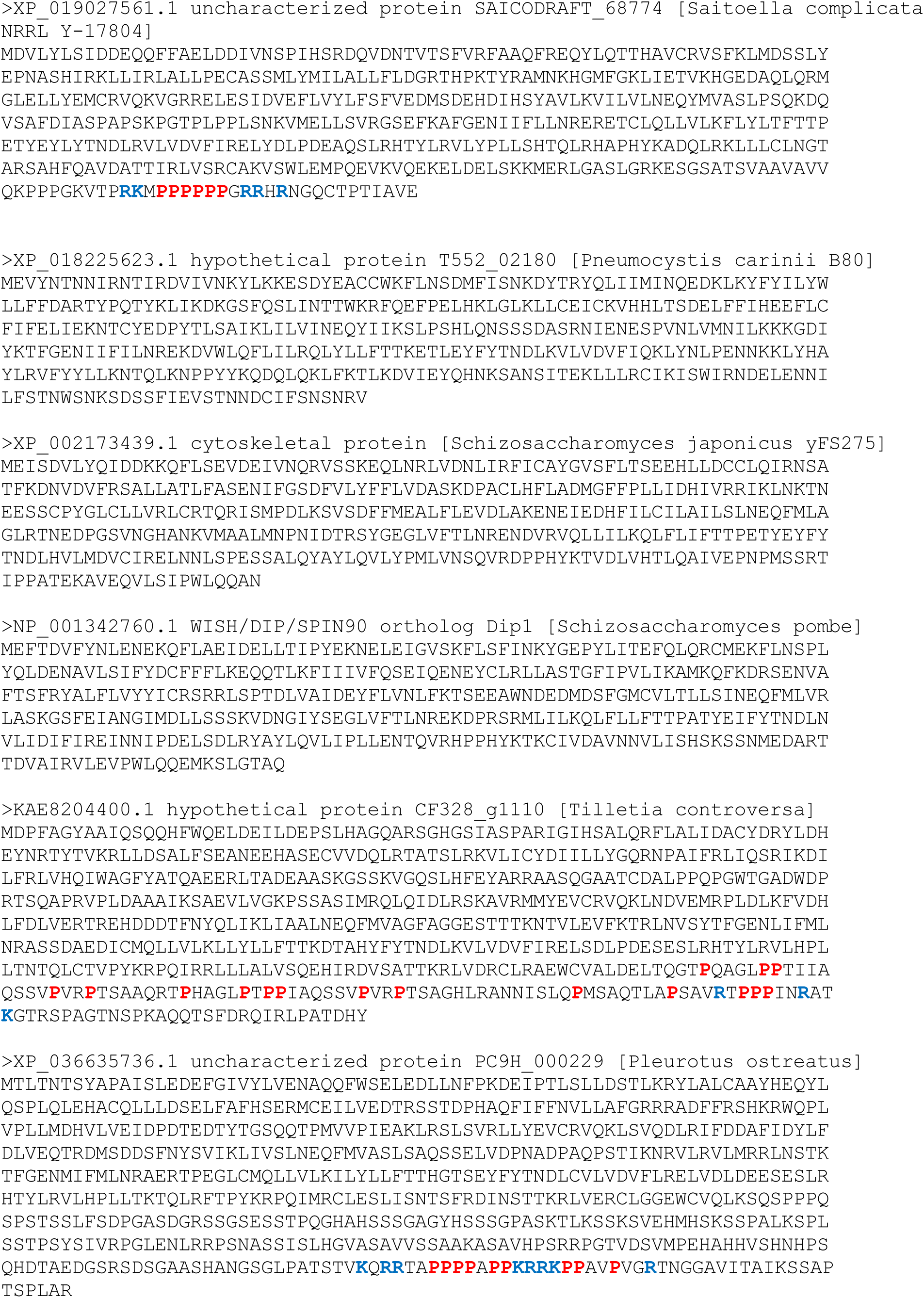
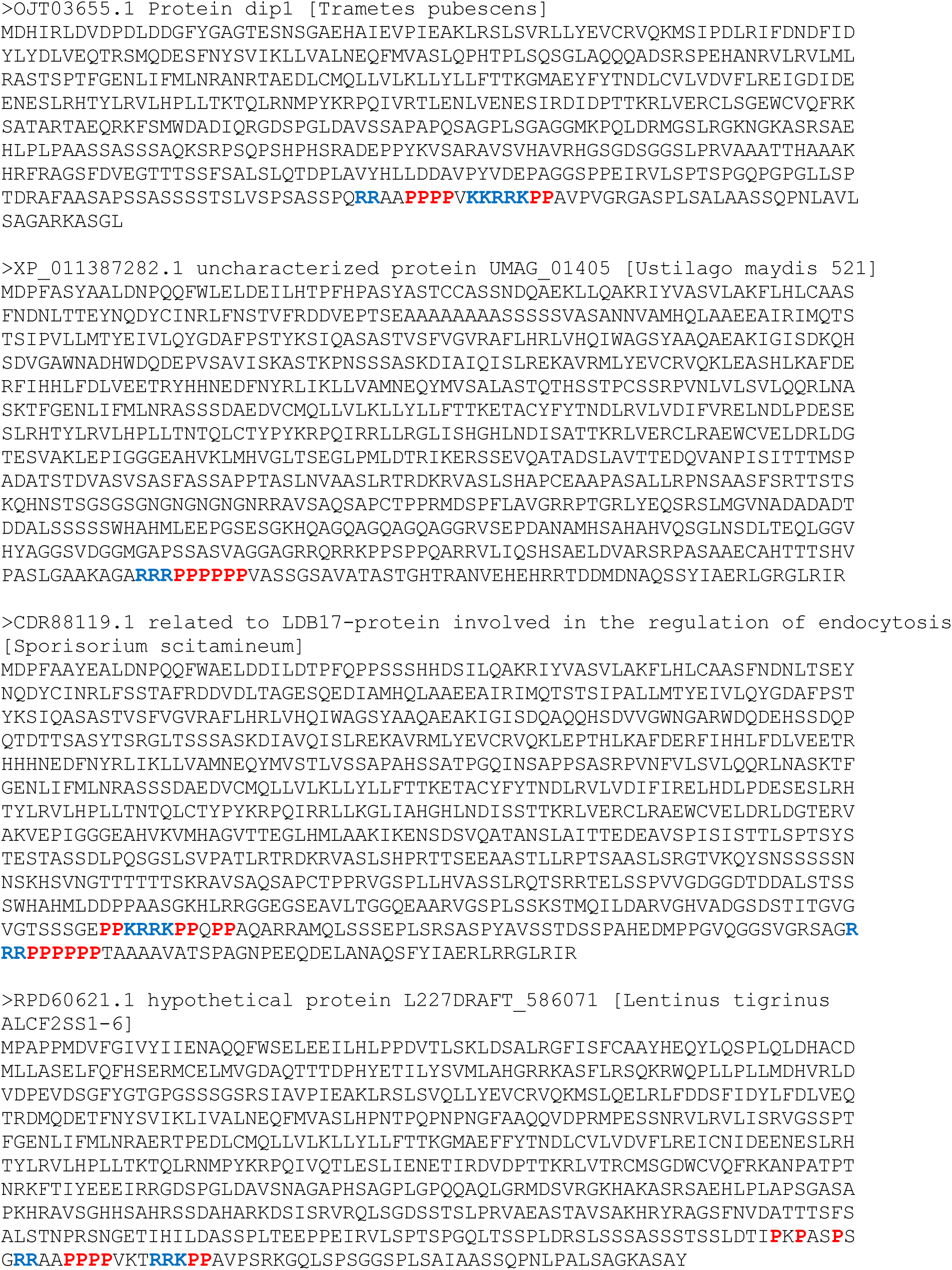
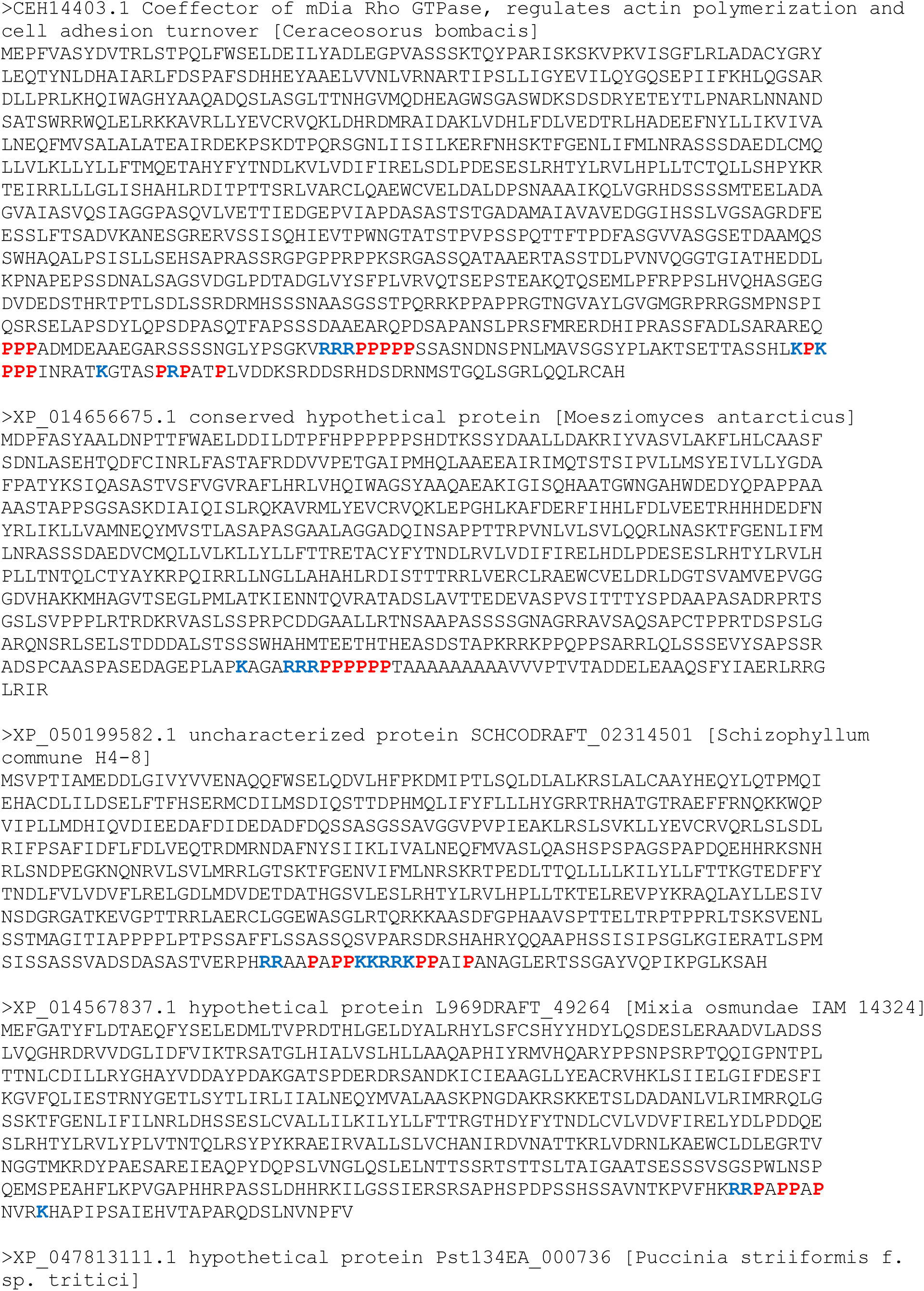
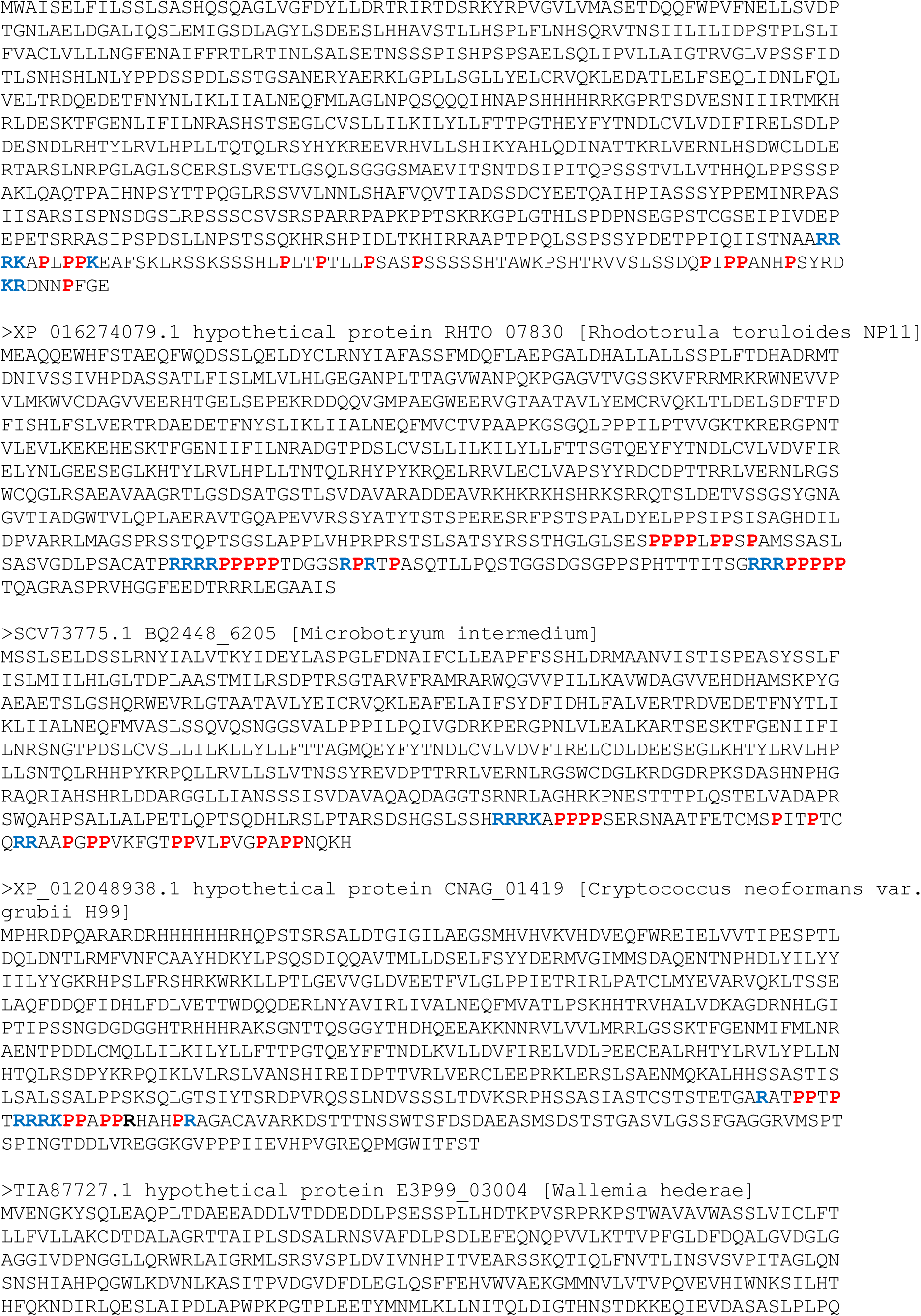
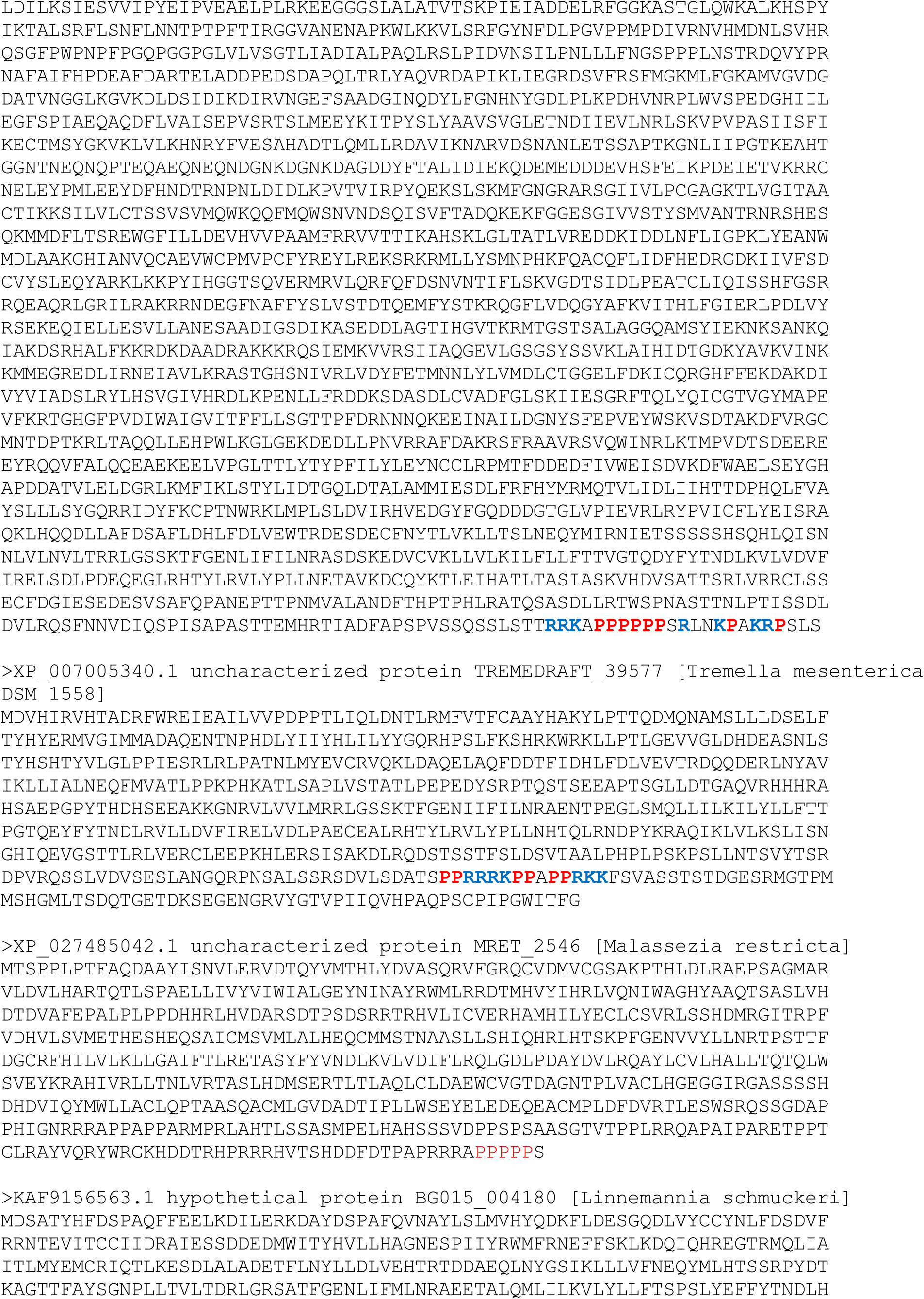
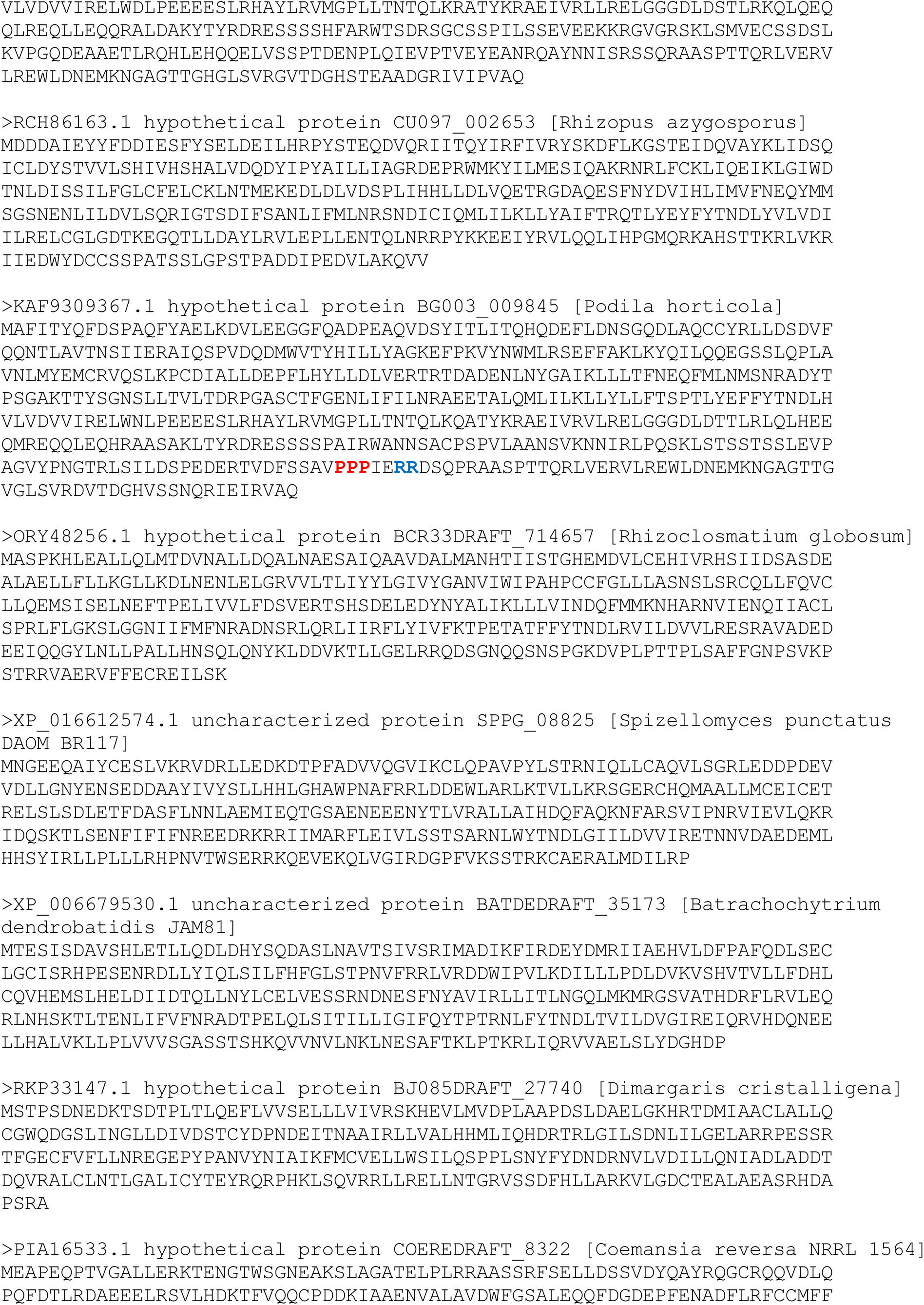
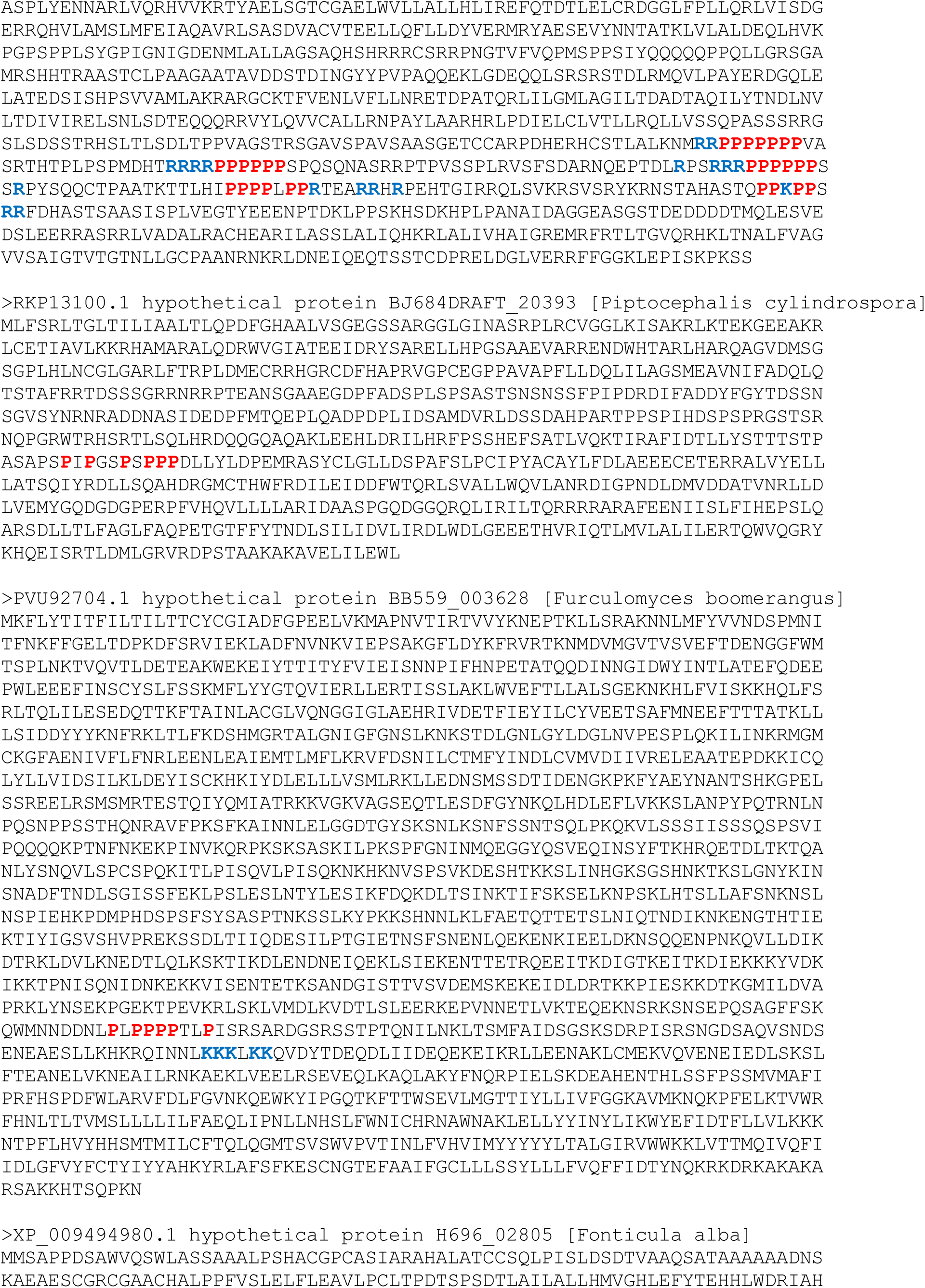
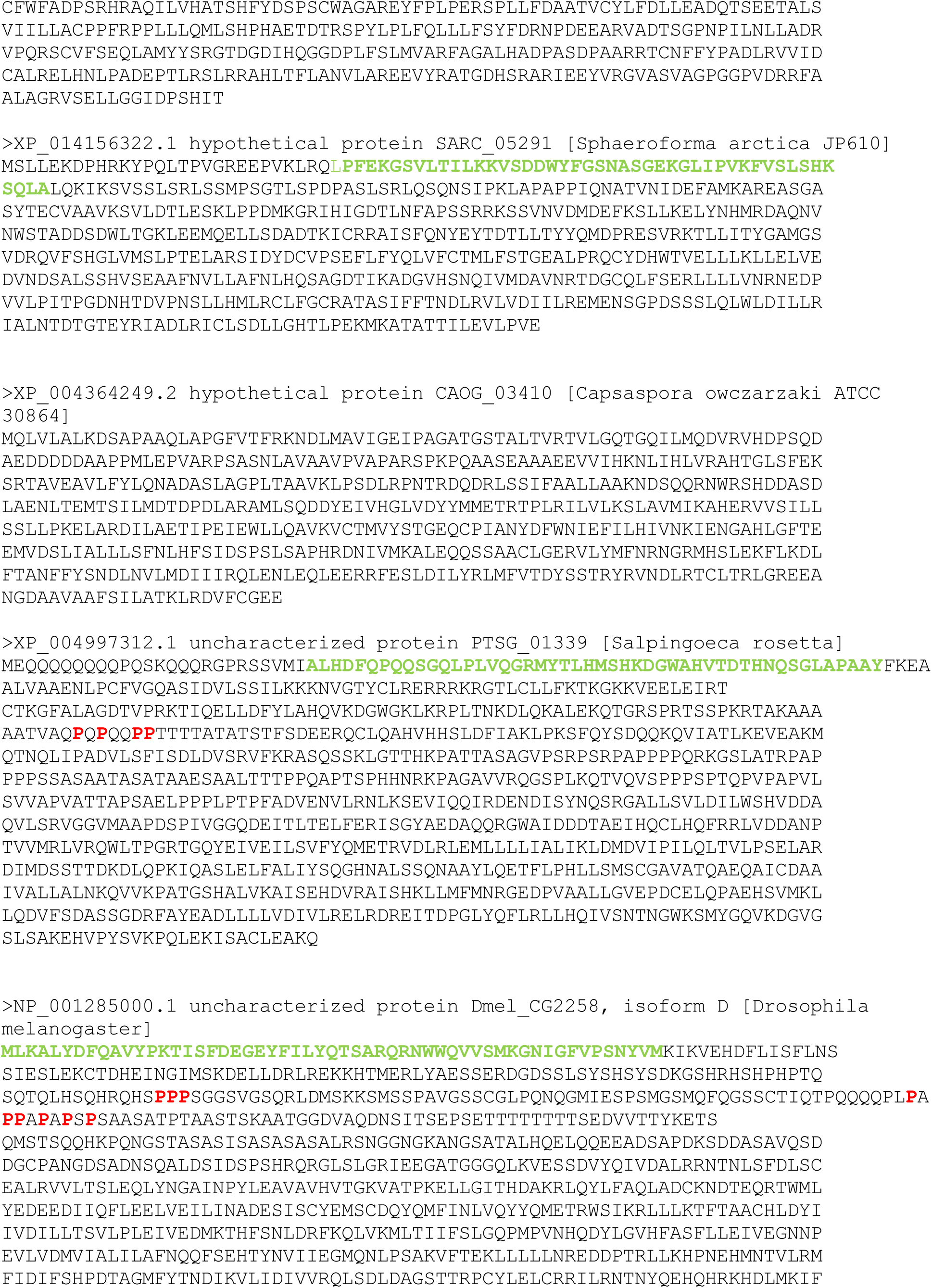
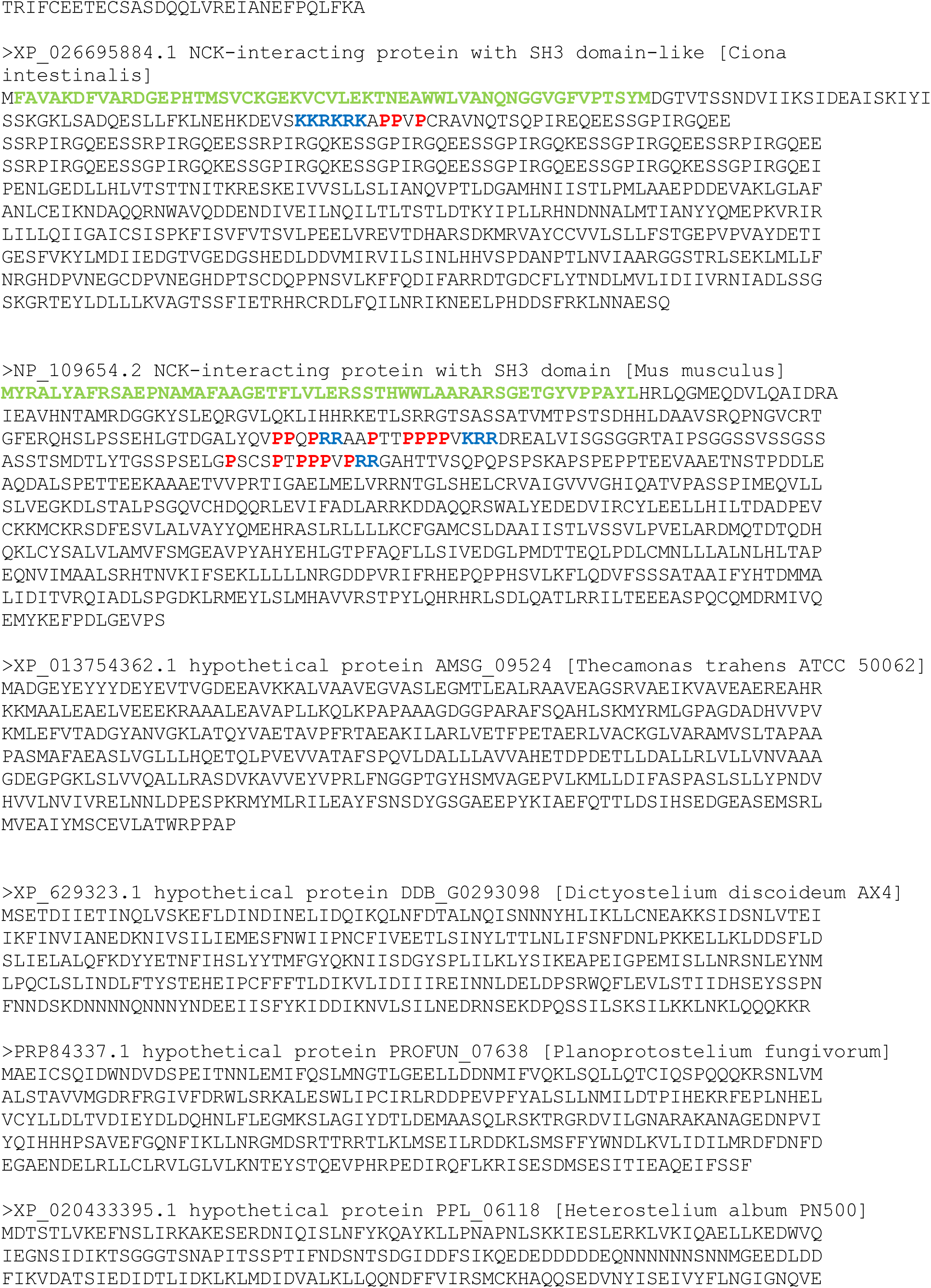
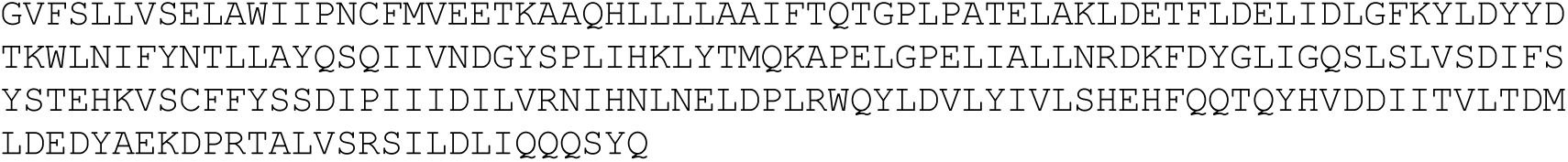

## Notes

### Competing Interest Statement

The authors have declared no competing interest.

